# Entorhinal grid-like codes and time-locked network dynamics track others navigating through space

**DOI:** 10.1101/2022.10.08.511403

**Authors:** Isabella C. Wagner, Luise P. Graichen, Boryana Todorova, Andre Lüttig, David B. Omer, Matthias Stangl, Claus Lamm

**Affiliations:** Social, Cognitive and Affective Neuroscience Unit, Department of Cognition, Emotion, and Methods in Psychology, Faculty of Psychology; Vienna Cognitive Science Hub; Department for Microbiology and Environmental Systems Science, University of Vienna, 1010 Vienna, Austria; Edmond and Lily Safra Center for Brain Sciences, The Hebrew University of Jerusalem, Givat Ram, 9190401 Jerusalem, Israel; Department of Psychiatry and Biobehavioral Sciences, Jane and Terry Semel Institute for Neuroscience and Human Behavior, University of California Los Angeles, Los Angeles, CA 90095, United States

**Keywords:** Grid cells, entorhinal cortex, socio-spatial navigation, brain networks, functional magnetic resonance imaging (fMRI)

## Abstract

Navigating through crowded, dynamically changing social environments requires the ability to keep track of other individuals. Grid cells in the entorhinal cortex are a central component of self-related navigation but whether they also track others’ movement is unclear. Here, we propose that entorhinal grid-like codes make an essential contribution to socio-spatial navigation. Sixty human participants underwent functional magnetic resonance imaging (fMRI) while observing and re-tracing different paths of a demonstrator that navigated a virtual reality environment. Results revealed that grid-like codes in the entorhinal cortex tracked the other individual navigating through space. Further, the activity of grid-like codes was time-locked to increases in co-activation and entorhinal-cortical connectivity that included the striatum, the hippocampus, parahippocampal and right posterior parietal cortices, altogether modulated by accuracy when subsequently re-tracing the paths. This suggests that network dynamics time-locked to entorhinal grid-cell-related activity might serve to distribute information about the ‘socio-spatial map’ throughout the brain.

## Introduction

Maneuvering a crowded sidewalk or coordinating with one’s team members to move towards the goal on a soccer field not only requires the planning of one’s own movement through space, but depends upon the ability to track the location of conspecifics. Such socio-spatial navigation was recently tied to neuronal processes similar to self-related spatial navigation. For instance, work using intracranial recordings from the human medial temporal lobe revealed representations that coded for environmental boundaries when participants or others moved through space (Stangl et al., 2021), and hippocampal social place cells were found to code for the location of others in bats (Omer et al., 2018) and rodents (Danjo et al., 2018). A central component of navigation is the integrity of grid cells in the entorhinal cortex that express periodic firing fields arranged along the vertices of regular hexagons, tessellating the environment and providing a spatial map (Hafting et al., 2005; Jacobs et al., 2013). In humans, it has been shown that the neural firing signature of grid cell populations can be measured non-invasively using fMRI, in the form of so-called “grid-like codes” (Doeller et al., 2010). Moreover, these grid-like codes were shown to support spatial (Doeller et al., 2010; Stangl et al., 2018) as well as mental self-related navigation (Bellmund et al., 2016; Horner et al., 2016), but whether they also track others’ movement through space is unclear. Here, we propose that entorhinal grid cells (and related grid-like codes) make an essential contribution to socio-spatial navigation in humans.

Similar to the joint actions of different spatial cell types such as place and grid cells (Bush et al., 2014; Fyhn et al., 2007; Moser et al., 2008; Zhang et al., 2013), flexible navigation relies on the orchestration of a set of regions including medial temporal, posterior-medial, parietal, striatal and prefrontal structures (Chersi and Burgess, 2015; Ekstrom et al., 2017; Epstein et al., 2017; Maguire et al., 1998; Patai and Spiers, 2021). Beyond the hippocampal-entorhinal circuit, the parahippocampal and retrosplenial cortices are regarded as key players involved in the visuospatial processing of scenes and their orientation in the broader spatial environment (Epstein, 2008; Epstein and Kanwisher, 1998; Maguire, 2001; Vann et al., 2009). The retrosplenial cortex specifically was proposed to translate information between allocentric (as supported by the hippocampus; Danjo, 2020) and egocentric reference frames (Ekstrom et al., 2017; Epstein, 2008; Vann et al., 2009), interacting with the (right) posterior parietal cortex to guide the visuospatial coordination of locomotion (Ciaramelli et al., 2010; McNaughton et al., 1994; Whitlock et al., 2008) and the coding of spatial routes (Nitz, 2006). Similar aspects of navigation such as goal-directed behavior (Chersi and Burgess, 2015) and route following (Hartley et al., 2003) were linked to dorsal striatal processes. Presumably, the dorsal striatum contributes to navigation via associative reinforcement and thus functions in parallel to the hippocampal system that rapidly encodes new experiences (Chersi and Burgess, 2015; Doeller et al., 2008; Iaria et al., 2003). Interactions between these two systems appear mediated by the prefrontal cortex (Doeller et al., 2008), which also regulates the top-down control of navigation via planning and goal tracking (Patai and Spiers, 2021). Since socio-spatial navigation also involves tracing others’ movement through space, it likely engages additional brain regions concerned with biological motion processing, such as the posterior superior temporal sulcus (Herrington et al., 2011; Sokolov et al., 2018). Altogether, this suggests a complex interplay of multiple brain regions, but it remains elusive how they interact with the putative “spatial map” hosted by entorhinal grid cells. We thus asked whether the activity of entorhinal grid-like codes is coupled to functional connectivity changes with medial temporal, parietal, striatal, and prefrontal areas and whether such network dynamics may explain differences in navigation performance.

To tackle these questions, we built upon the design of previous animal work that investigated social place cells in bats (Omer et al., 2018) and translated it into the human context. Sixty healthy participants underwent functional magnetic resonance imaging (fMRI). Our modified navigation task projected them into a first-person perspective within a virtual reality (VR) environment while they were asked to observe different paths of a demonstrator (**Figure 1AB**). The paths had to be held in memory and needed to be re-traced after a short delay. Crucially, participants were positioned at a fixed viewpoint during observation, allowing us to dissociate potential grid-like codes between other- and self-related movements. Behavioral performance was quantified as cumulative distance error in virtual meters (vm) indicating the deviation from the demonstrator’s paths. We reasoned that if underlying grid cells supported the tracking of others, we should observe increased grid-like codes in the entorhinal cortex during observation of the demonstrator moving through VR-space. Furthermore, we expected entorhinal grid-like codes to be dovetailed by entorhinal-cortical connectivity changes with regions typically involved in spatial navigation and visuospatial processing, altogether modulated by behavioral performance.

**Figure 1.**
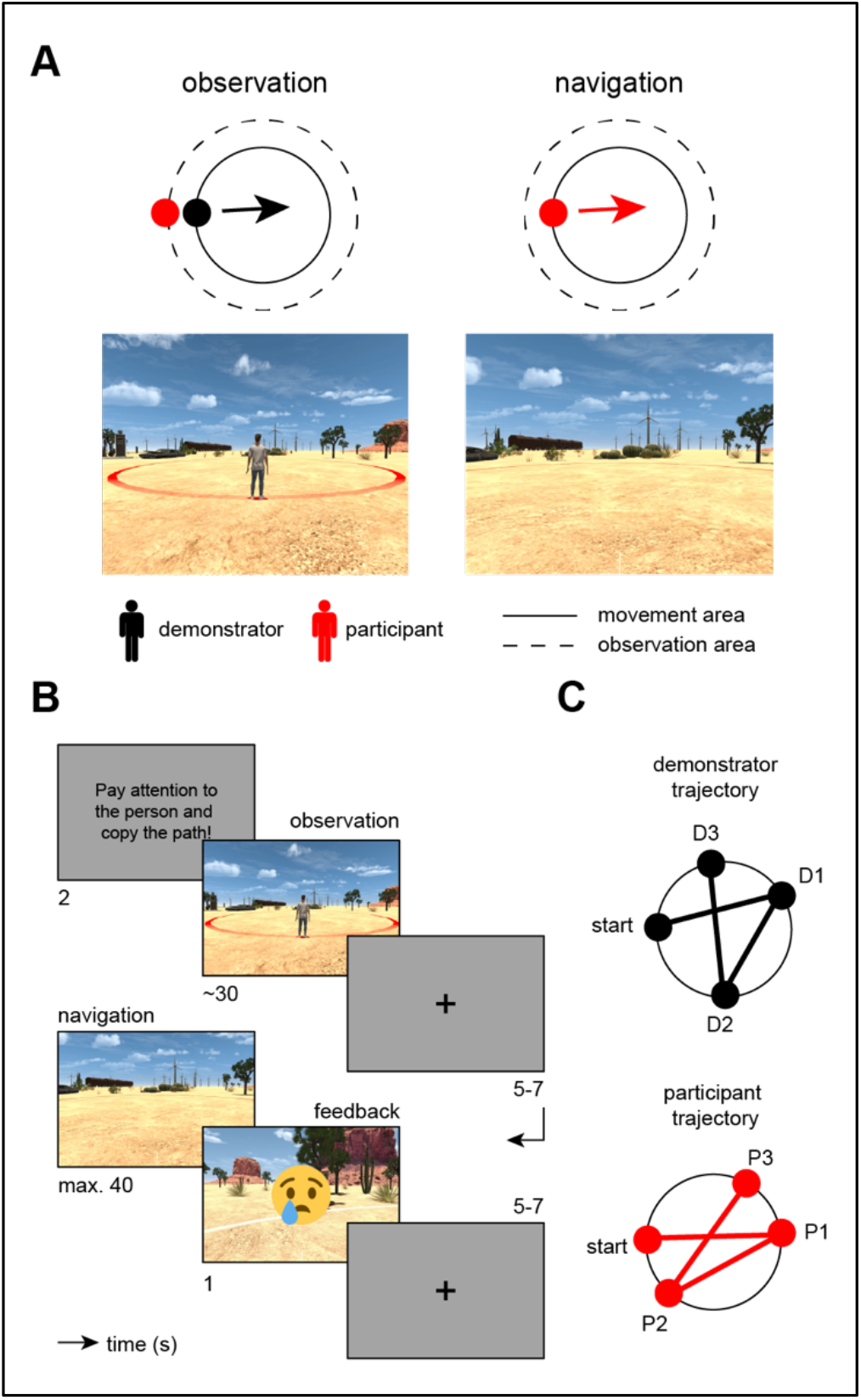
Modified navigation task. **(A)** The task projected participants into a first-person perspective within a virtual reality (VR) environment while they were asked to observe and subsequently re-trace the paths of a demonstrator (screenshots). The movement area was marked through a red circle on the sandy desert plane (solid circle as indicated in bird’s eye view, upper panels), and was surrounded by an observation area (dashed line in bird’s eye view). During observation (left panel), participants were placed directly behind the demonstrator’s starting point, were not able to move, and viewed the demonstrator walking through the circular arena. During navigation (right panel), the demonstrator disappeared and the participant was virtually projected onto the same starting position to re-trace the previously observed path (arrow exemplifies participant movement). **(B)** Timeline of one example trial during the modified navigation task (s, seconds). **(C)** Example path of demonstrator (D) and participant (P) obtained from observation and navigation periods, respectively. Each trial comprised three random path segments (D/P1-3) that each started and ended at the edge of the movement area (i.e., participants observed/walked paths from one edge to the other and were not able to stop and turn within the movement area).

## Results

### Participants are successfully able to re-trace the demonstrator’s paths

Participants completed a single MRI session that started out with an initial task familiarization period, followed by four runs of the modified navigation task, each involving different paths. Across the four task runs, participants reached an average cumulative distance error of 17.57 ± 0.54 vm (mean ± standard error of the mean, s.e.m.; calculated as the average distance between the demonstrator’s and the participant’s endpoints across the three different segments of a given path, **Figure 1C**), significantly below the threshold of 20 vm which determined the feedback that participants received [i.e., happy emoji, cumulative distance error =< 20 vm; sad emoji, cumulative distance error > 20 vm; one-sample *t*-test, *N* = 58 (2 participants were excluded from this and all following analyses, **Materials and Methods**); *t*(57) = - 4.49, Cohen’s *d* = -0.59, 95% confidence interval (CI) = [16.5, 18.7], *p*_two-tailed_ < 0.0001]. Performance was stable throughout the experiment (**Figure S1**), showing that participants were successfully able to re-trace the demonstrator’s paths, and providing us with a solid basis for investigating grid-like codes during both observation and navigation.

We next took a closer look at navigation performance on individual segments of a given path (each path consisted of three segments). We found that across-run performance was significantly better when participants re-traced the first path segment (6.86 ± 0.16 vm) compared to the second (23.3 ± 0.32 vm) or third (22.64 ± 0.37 vm; as indicated by the path segment-wise distance error; **Figure S1**). This pattern was evident in each of the four runs (**Figure S1**) and was likely due to the fact that the observer’s viewpoint was directly fixed behind the demonstrator’s starting position at the beginning of each trial.

### Observation is associated with increased activation in the hippocampus and striatum

We then turned towards the fMRI data and started out by investigating changes in brain activation when participants tracked and re-traced the paths of others, irrespective of potential grid-like codes. We hypothesized largely overlapping activation profiles during both observation and navigation periods, involving regions typically engaged in spatial navigation and visuospatial processing, such as medial temporal, posterior-medial, parietal, striatal, and prefrontal areas (Chersi and Burgess, 2015; Ekstrom et al., 2017; Epstein et al., 2017; Maguire et al., 1998; Patai and Spiers, 2021). Additionally, we expected that observation of the demonstrator’s paths would be associated with increased activation in brain areas associated with biological motion processing, including the posterior superior temporal sulcus (Herrington et al., 2011; Sokolov et al., 2018).

While participants observed the demonstrator’s paths, a set of regions showed increased activation, including a large cluster centered on the bilateral hippocampus and adjacent structures of the medial temporal lobe (MTL), extending towards the anterior temporal pole, the caudate nucleus, pre- and post-central gyri, as well as the ventromedial prefrontal and posterior cingulate cortices (one-sample *t*-test, *N* = 58, contrast observation > navigation; **Figure 2A, Table S1**). Results further included stronger bilateral activation in lateral occipital areas, the fusiform gyrus, and the posterior superior temporal sulcus.

**Figure 2.**
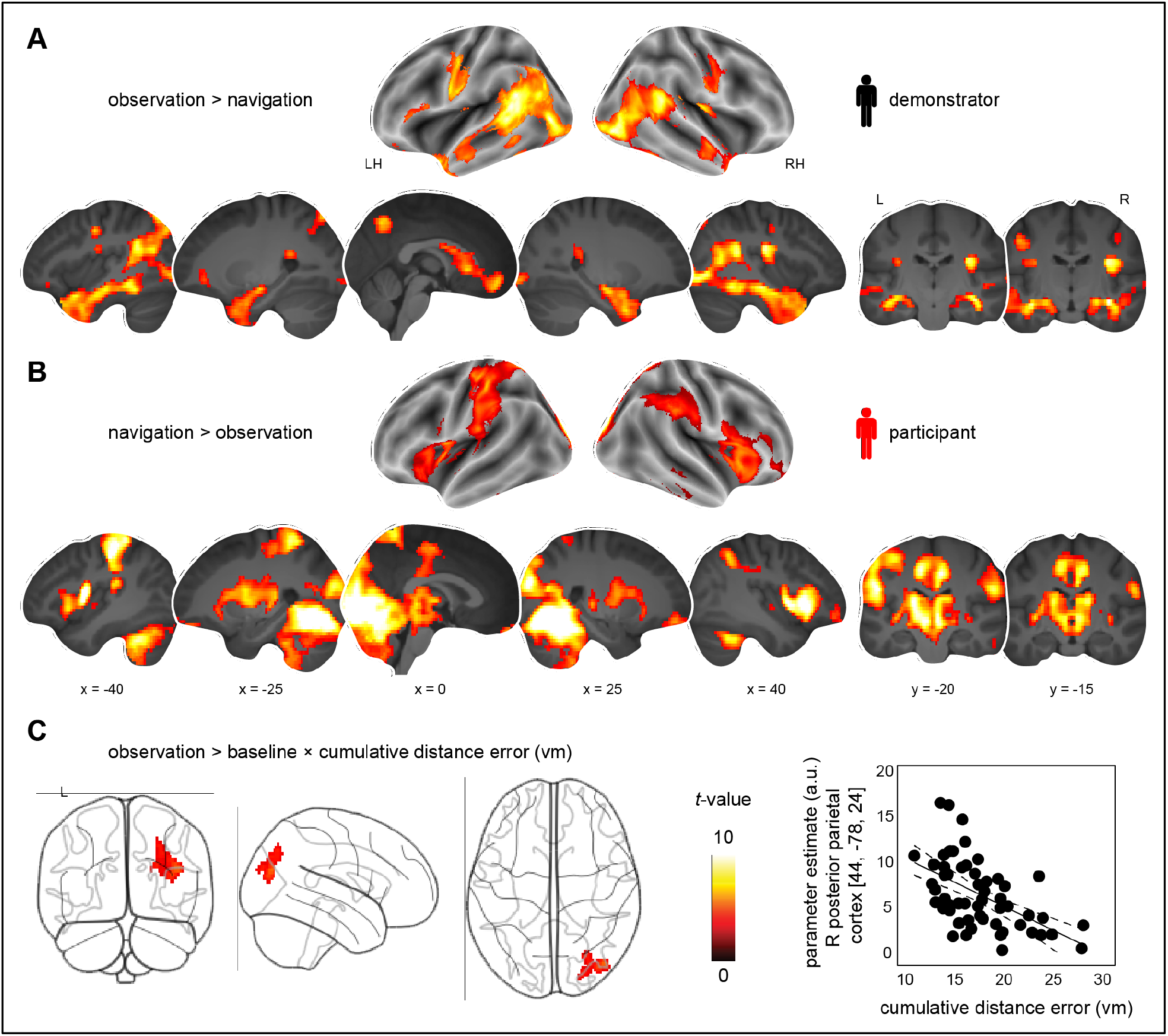
Brain activation profiles during observation and navigation, and association with performance. Brain activation **(A)** during observation (compared to navigation) periods, and **(B)** vice versa (**Table S1, Figure S2** for contrasts of each condition against the implicit fixation baseline). **(C)** Brain activation during observation (compared to implicit baseline) and association with performance across participants (indexed by the average cumulative distance error in virtual meters, vm). The scatter plot shows the relationship between the change in parameter estimates (arbitrary units, a.u.) extracted from the significant cluster and the cumulative distance error (vm; **Results S1**). Given the clear inferential circularity, we would like to highlight that this plot serves visualization purposes only, solely illustrating the direction of the brain-behavior relationship. All results are shown at *p* < 0.05 FWE-corrected at cluster level (cluster-defining threshold of *p* < 0.001).

When participants re-traced the previously observed paths during navigation periods, an activation profile with strongest response in the bilateral occipital cortex, including the fusiform gyrus and the right temporo-parietal junction emerged (one-sample *t*-test, *N* = 58, contrast navigation > observation; **Figure 2B, Table S1**). Activation was also increased in the bilateral caudate nucleus and putamen, the insula, as well as the adjacent inferior prefrontal cortex. Notably, the general effect of navigation (contrast navigation > baseline) was also associated with increased activation in the hippocampus and entorhinal cortex (**Figure S2, Table S1**).

Altogether, observing the demonstrator’s paths rather than re-tracing them was associated with stronger brain activity in a set of regions that included the hippocampus and adjacent MTL structures, the striatum, anterior and posterior midline regions, as well as superior and inferior temporal areas.

### Increased activation in the right posterior parietal cortex during observation is associated with better performance

To investigate whether brain activation during observation and navigation scaled with individual performance, we went on to test the cross-participant relationship between whole-brain activity during observation/navigation periods and the average cumulative distance error when re-tracing the demonstrator’s paths. Results showed an activation increase within the right posterior parietal cortex (*x* = 44, *y* = -78, *z* = 24, *z*-value = 4.68, 297 voxels) during observation that negatively correlated with the individual cumulative distance error [linear regression, *N* = 58, contrast observation > baseline, cumulative distance error added as a covariate of interest; *p* < 0.05 family-wise-error (FWE) corrected at cluster level using a cluster-defining threshold of *p* < 0.001, cluster size = 80 voxels; **Figure 2C**]. In other words, stronger activation in this region was coupled to better subsequent performance as participants observed the demonstrator’s paths (and see **Results S1** for additional analysis). We did not find a significant relationship between behavioral performance and brain activation during navigation periods.

### Entorhinal grid-like codes when observing the demonstrator’s paths

Central to our analyses was the question whether grid-like codes supported the tracking of others. We were motivated by findings of place cells in animals that were recently shown to signal the location of conspecifics (Danjo et al., 2018; Omer et al., 2018), as well as by previous work probing entorhinal grid cell activity (or grid-like codes) during spatial (and mental) self-navigation in both animals (Hafting et al., 2005) and humans (Bellmund et al., 2016; Doeller et al., 2010; Horner et al., 2016; Stangl et al., 2018). Hence, we expected to find a significant hexadirectional modulation (i.e., a 6-fold periodicity) of the fMRI signal in the entorhinal cortex during observation and possibly also during navigation periods (**Figure 3A**).

**Figure 3.**
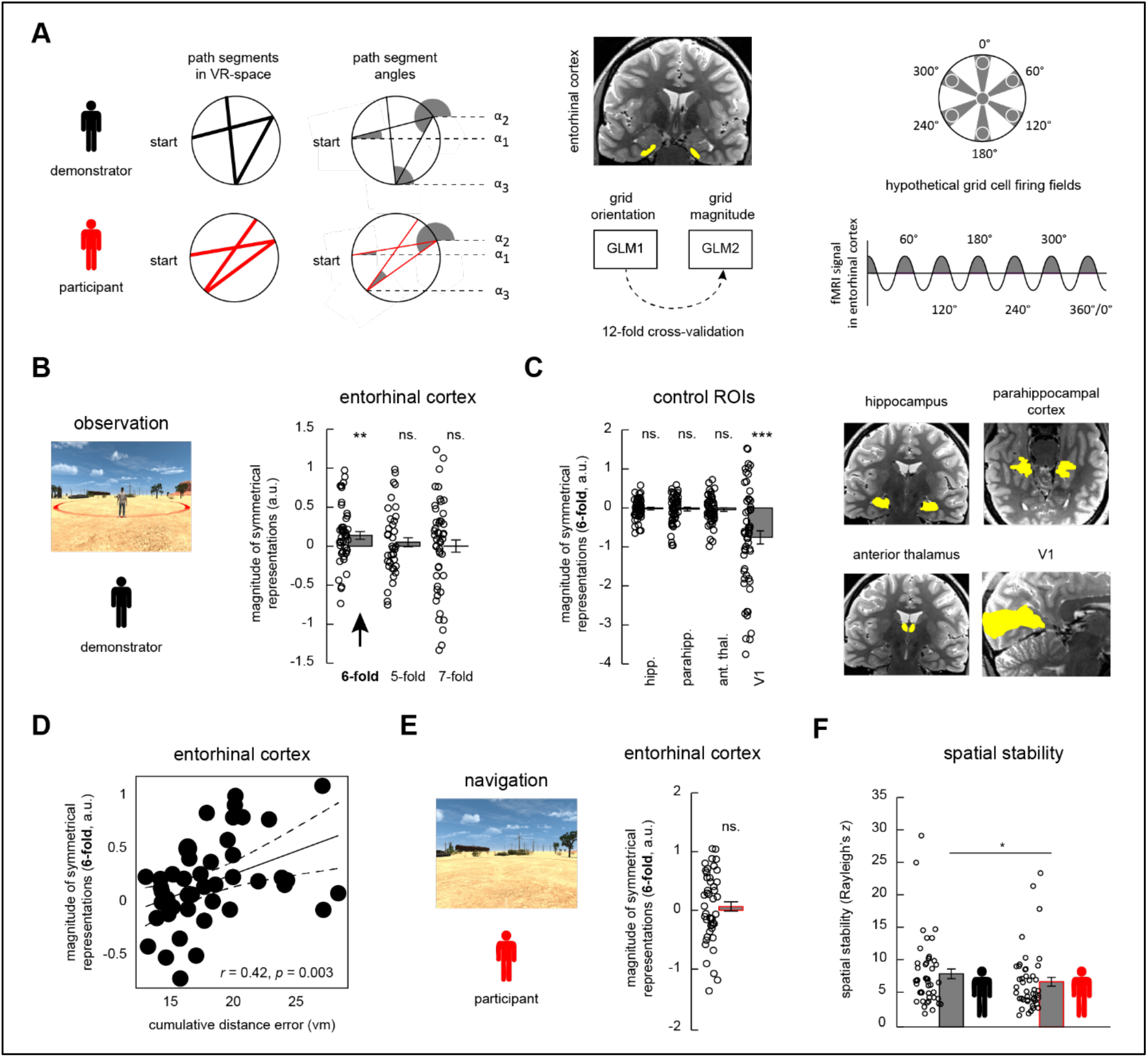
Grid-like codes during observation. **(A, left panel)** We calculated path-wise trajectory angles α_1_-α_3_ referenced to an arbitrary point on the VR desert plane (dashed lines, angles indicated in grey). **(A, middle panel)** Entorhinal cortex region-of-interest (ROI, in yellow) projected onto the T2-weighted structural scan of one participant. Grid orientations were estimated and tested by employing a 12-fold cross-validation (CV) regime. **(A, right panel)** Hexadirectional modulation of hypothetical grid cell firing fields (indicated as white circles). The fMRI signal in the entorhinal cortex is expected to be increased whenever participants observe the demonstrator walking (or whenever participants walk) aligned with their internal grid orientation (highlighted in grey). **(B)** Magnitude of symmetrical representations (6-, 5-, and 7-fold periodicities) in the entorhinal cortex during observation periods (arbitrary units, a.u.; **Results S2**). **(C)** Magnitude of observation-related symmetrical representations (6-fold periodicity) in control ROIs. **(D)** Correlation of grid magnitudes (6-fold periodicity) and cumulative distance error across participants (vm). **(E)** No significant entorhinal symmetrical representations (6-fold periodicity) during navigation periods (**Results S3**). **(F)** Spatial stability of entorhinal grid-like codes. *** *p* < 0.0001, ** *p* < 0.01, * *p* < 0.05; error bars reflect the standard error of the mean, s.e.m.

To test this, we split the fMRI data of each task run into independent sets and estimated/tested individual grid orientations using a 12-fold cross-validation regime (for a detailed description, please see **Materials and Methods**). We used 11 trials (each trial included one observation and navigation period) to estimate individual grid orientations based on the demonstrator’s (or the participant’s) movement trajectories through VR-space using a General Linear Model (GLM1; note that grid orientations were estimated separately for observation and navigation periods). The grid orientations’ fit was then tested on the path segments of the remaining trial, which served to quantify the magnitude of grid-like codes in the left-out data set (GLM2). This procedure was repeated for each cross-validation fold and the resulting grid magnitudes were averaged across the different iterations. Note that grid analyses were based on a reduced participant sample (entorhinal cortex: automatic segmentation, *N* = 49; manual segmentation, *N* = 51; control regions: *N* = 58; **Materials and Methods**).

We found significant grid-like codes in the entorhinal cortex as participants were observing the demonstrator’s paths. This effect was only present for the 6-fold symmetrical model [mean grid magnitude (arbitrary units, a.u.) ± s.e.m., 0.158 ± 0.06; one-sample *t*-test, *N* = 47 (excluding 2 outliers); *t*(46) = 2.73, *d* = 0.4, 95% CI = [0.04, 0.3], *p*_one-tailed_ = 0.005, Bonferroni-corrected for multiple comparisons using a threshold of α_Bonferroni_ = 0.05/ a total of 6 entorhinal and control ROIs = 0.008; **Figure 3B**; results for this and all following analyses of this section remained stable when using the full data set of *N* = 49 and not excluding any outliers, see also **Table S2**]. Different symmetrical models such as 5- or 7-fold signal periodicities did not yield significant results [separate one-sample *t*-tests; 5-fold: *N* = 42 (excluding 7 outliers); *p*_one-tailed_ = 0.245; 7-fold: *N* = 48 (excluding 1 outlier), *p*_one-tailed_ = 0.489; **Figure 3B**]. The same pattern of findings emerged when we delineated the entorhinal cortex using manual segmentation (**Results S2, Figure S3**), and when using a different, less nested cross-validation regime that split each task run into four data parts (**Results S2**). Exploratory follow-up analysis showed that grid-like codes during observation appeared predominantly left lateralized (**Results S2**).

To test whether grid-like codes were also present in other areas, we chose several control ROIs known to be involved in spatial navigation and visuo-spatial processing but for which no grid-like codes have been reported so far (including the hippocampus, parahippocampal cortex, anterior thalamus, and the primary visual cortex/V1; **Figure 3C, Materials and Methods**). There were no significant grid-like codes in any of the control ROIs [separate one-sample *t*-tests; hippocampus: *N* = 56 (excluding 2 outliers), *p*_one-tailed_ = 0.271; parahippocampal cortex: *N* = 55 (excluding 3 outliers), *p*_one-tailed_ = 0.264; anterior thalamus: *N* = 56 (excluding 2 outliers), *p*_one-tailed_ = 0.159; V1: *N* = 58 (full data set, no outliers excluded), significantly negative grid magnitude, *d* = -0.59, 95% CI = [-1.1, -0.4], *p*_one-tailed_ < 0.0001; **Figure 3C**; and see **Table S2** for virtually identical results when including the full data set]. We further explored the finding of negative grid-like codes in V1 during observation with additional analyses and compared to previous work (**Results S2**). In summary, we found significantly increased grid-like signals in the entorhinal cortex, but not in any of the control regions, when participants were observing and putatively encoding the demonstrator’s paths.

### Entorhinal grid-like codes during observation are stronger at lower performance

Previous studies highlighted a significant association between entorhinal grid-like codes during navigation and behavioral performance (Doeller et al., 2010). We thus reasoned that variations in observation-based grid-like codes (as indexed by grid magnitudes) should be coupled to individual differences in the average cumulative distance error during navigation periods. Interestingly, results yielded a significant positive association such that increased observation-based grid magnitudes were coupled to larger cumulative distance errors (*N* = 47, same sample as for main analysis above: *r*_*Pearson*_ = 0.42, 95% CI = [0.15,0.63], *p*_two-tailed_ = 0.003; **Figure 3D**).

### No significant entorhinal grid-like codes when re-tracing the demonstrator’s paths

In analogy to previous work that reported entorhinal grid cell activity, or grid-like codes, during spatial (and mental) self-navigation (Bellmund et al., 2016; Doeller et al., 2010; Hafting et al., 2005; Horner et al., 2016; Jacobs et al., 2013; Stangl et al., 2018), we expected to replicate this finding in our data set and hypothesized significant grid-like codes in the entorhinal cortex as participants were re-tracing the demonstrator’s paths during navigation periods (**Materials and Methods**). Surprisingly, results did not show significantly increased grid-like codes in the entorhinal cortex [separate one-sample *t*-tests for all following analyses; *N* = 45 (excluding 4 outliers), *p*_one-tailed_ = 0.233; **Figure 3E**; as above, results for this and all following analyses of this section remained the same when using the full data set, **Table S3**], also not when manually delineating the region, when using a reduced data set (**Results S3**), when using a different cross-validation regime (**Results S3**), or when testing for grid-like codes in the left and right hemisphere separately (**Results S3**). When investigating grid-like codes in the control ROIs, we found significantly increased grid magnitudes in the anterior thalamus during navigation but not in any of the other regions (**Results S3, Figure S3**).

We also explored whether grid orientations during observation periods served as spatial reference frames when participants re-traced the demonstrator’s paths during navigation. If this was the case, we should find matching grid orientations between the two conditions, leading to significantly increased grid magnitudes when testing grid orientations obtained from observation on navigation periods. However, results did not yield any significant results, indicating grid-like codes during observation but neither the same nor differently-oriented grid-like codes during navigation periods (**Results S3, Figure S3**).

### Control analyses

#### Unbiased estimation of grid orientations

We performed several control analyses to verify that our results were actually related to entorhinal grid-like codes that coded for the demonstrator’s path in space. For instance, the estimation of individual grid orientations can be biased by the distance walked along different directions in space (in other words, it would not be possible to estimate individual grid orientations for a specific direction if the participants never walked in that direction). To circumvent this issue, we designed our task such that each segment of an individual path was oriented along one of 36 directions that could be divided into 6 directional bins, spanning the 360°-VR-space with 10° angular resolution (Doeller et al., 2010). We then generated the demonstrator’s paths (i.e., the entire trajectories that consisted of three path segments each) to maximize the distance walked in each of the directional bins and could thus avoid any biases in the estimation of individual grid orientations during observation periods (**Materials and Methods, Figure S4**).

Since participants were not always able to perfectly re-trace the demonstrator’s paths, we also checked whether there was a difference in the participant’s distance walked across the 6 directional bins. As above, there was no significant difference in the average distance walked suggesting that we were able to avoid any biases in grid orientation estimation during navigation periods (**Figure S4**).

#### Higher spatial stability of entorhinal grid-like codes during observation

Grid-like codes might be affected by variations in spatial and temporal signal stability (Stangl et al., 2018). In the case of spatial instability, estimated grid orientations are assumed to vary across the different voxels within the ROI, resulting in more variable mean grid orientations and an overall decrease in grid magnitude. To test whether differences in spatial stability between observation and navigation conditions contributed to our results, we calculated individual voxel-wise grid orientations within the bilateral entorhinal cortex ROI (**Materials and Methods**).

Results revealed that the spatial stability [quantified as Rayleigh’s *z* that describes non-uniformity of circular data; i.e., data clustering towards a specific (grid) orientation] was significantly higher for grid-like codes during observation compared to navigation periods [paired-sample *t*-test, *N* = 44, participant sample from which both observation- and navigation-based grid values were available, *t*(43) = 2.04, *d* = 0.28, 95% CI = [0.01, 2.4], *p*_two-tailed_ = 0.048; **Figure 3F**]. Voxel-wise entorhinal grid orientations thus varied more when participants re-traced the demonstrator’s paths, potentially contributing to the lack of significant grid magnitudes during navigation. There was no significant difference in the temporal stability of voxel-wise entorhinal grid magnitudes between observation and navigation periods over time (i.e., no difference in grid orientations in the estimation and test data sets, quantified as % voxels with same/different orientations across time; **Materials and Methods, Figure S4**).

#### No effects of eye movements on grid-like codes

To account for the potential impact of eye movements on grid-like codes (Julian et al., 2018; Killian et al., 2012; Nau et al., 2018; Staudigl et al., 2018), we recorded eye gaze during the modified navigation task and leveraged the saccade directions of each participant during observation and navigation periods (note that this analysis included a subset of 37 participants from which both eye-tracking and entorhinal fMRI data were available). We repeated our initial grid analysis but now modelled eye gaze directions (i.e., the angle between successive saccades with respect to an arbitrary reference point on the computer screen) instead of movement trajectories in VR-space (**Results S4**). Findings did not reveal a significant increase in entorhinal saccade-based grid magnitudes when testing for a 6-fold symmetrical model during observation (or navigation) periods (separate one-sample *t*-tests, all *p*_two-tailed_ > 0.05; **Results S4, Figure S4**), altogether suggesting that our result of significant grid-like codes in the entorhinal cortex when observing the demonstrator’s paths was based on spatial information rather than eye movements.

We also verified whether participants were actually following the demonstrator with their eye gaze during observation by defining an area-of-interest (AOI) for each path segment. AOIs were defined by the two-dimensional coordinates of a given path trajectory on the computer screen as well as by the demonstrator’s height, and we calculated the percentage of eye movements that were located within the AOI boundaries. Indeed, the majority of eye movements were located within AOIs, emphasizing that participants’ viewing behavior was related to the observation of the demonstrator (**Results S4**).

### Striatal activation increases are time-locked to entorhinal grid-like codes during observation and negatively scale with performance

We next asked whether the activity of entorhinal grid-like codes during observation was paralleled by activation changes of regions typically involved in spatial navigation and visuospatial processing (e.g., medial temporal, parietal, striatal, and prefrontal structures; Chersi and Burgess, 2015; Ekstrom et al., 2017; Epstein et al., 2017; Maguire et al., 1998; Patai and Spiers, 2021), and whether this would be linked to behavioral performance. Similar to above, we modelled observation and navigation periods based on the individual path segments but now added the participant- and path segment-specific grid magnitudes (obtained from the previous grid code analysis) as parametric modulators (GLM3; grid magnitudes were obtained from GLM2 and were then averaged across the 12 cross-validation folds, **Figure 4A, Materials and Methods**). We then performed group comparisons to estimate voxel-wise activation changes that scaled with participant- and path segment-specific grid magnitudes (i.e., contrasting the parametric modulation regressor that captured fluctuations in entorhinal grid magnitude during observation/navigation against the implicit baseline).

**Figure 4.**
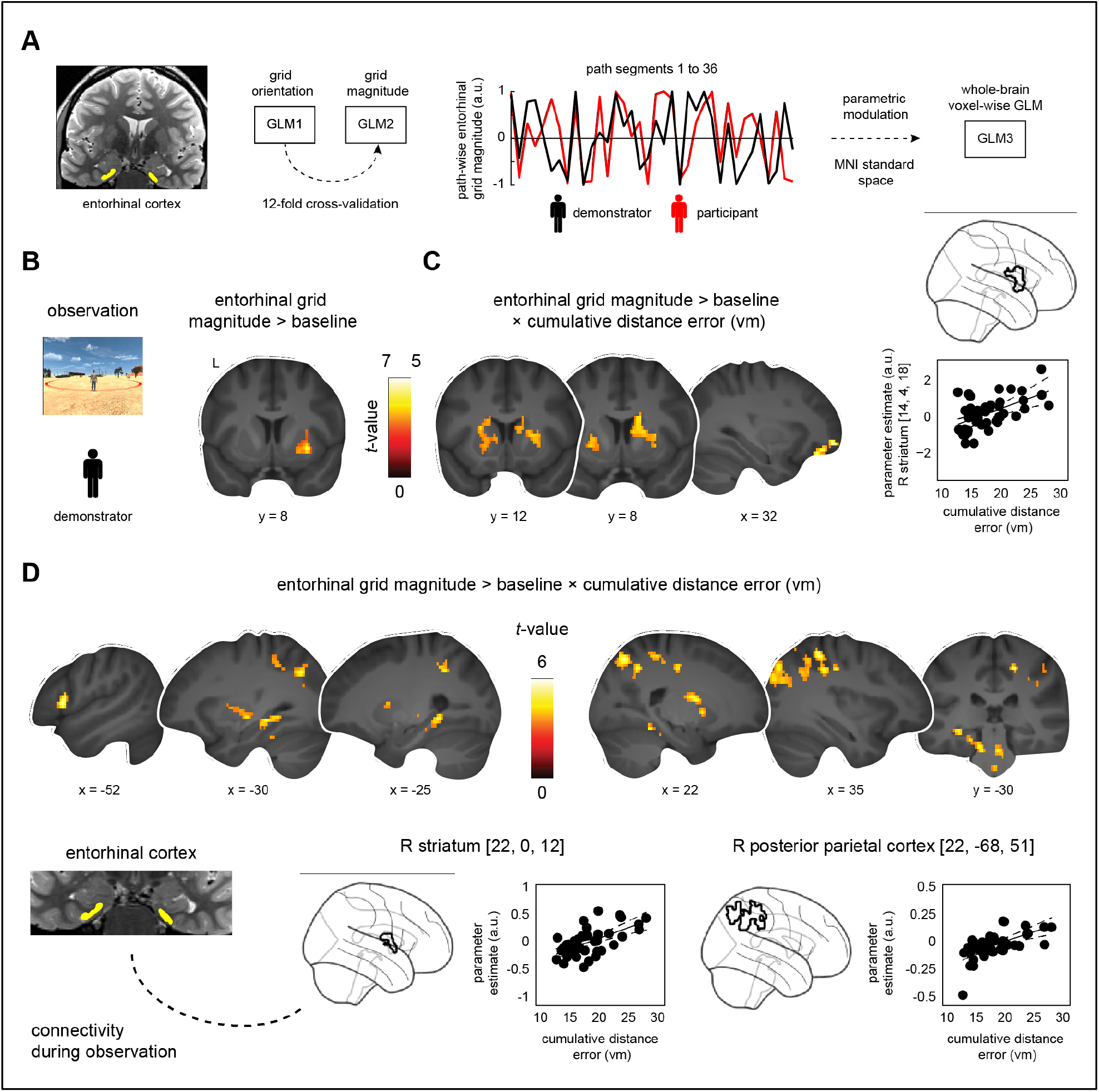
Brain activation profiles and connectivity changes time-locked to entorhinal grid-like codes during observation, and association with performance. **(A)** Grid orientations and magnitudes were estimated and tested on independent data sets (GLM1 and GLM2), focusing on data within the entorhinal cortex (yellow). This yielded participant- and path segment-specific grid magnitudes for each of the 36 path segments per task run. Segment-wise grid magnitudes were then used to parametrically modulate observation/navigation events in a third, independent analysis (GLM3) that tested for voxel-wise brain activation/connectivity changes time-locked to fluctuations in entorhinal grid magnitude. **(B)** Increased brain activation during observation periods (compared to the implicit baseline), positively scaling with entorhinal grid magnitudes. **(C)** Same contrast associated with behavioral performance. **(D)** Entorhinal connectivity changes during observation periods, time-locked to increased entorhinal grid magnitudes and associated with behavioral performance (**Table S4-5, Figure S5-6**). **(C-D)** The scatter plots show the relationships between the changes in parameter estimates (arbitrary units, a.u.) extracted from the respective clusters (black outlines in glass brains) and the cumulative distance error (vm). Given the clear inferential circularity, we would like to highlight that these plots serve visualization purposes only, solely illustrating the direction of the brain-behavior relationship. Results are shown at *p* < 0.05 FWE-corrected at cluster level [cluster-defining threshold of **(A**,**D)** *p* < 0.001 or **(C)** *p* < 0.005].

During observation periods, we found significantly increased activation in the right putamen and caudate nucleus (*x* = 30, *y* = 8, *z* = -6, *z*-value = 4.96, 102 voxels) that positively scaled with individual entorhinal grid magnitudes as participants were observing the demonstrator’s paths (one-sample *t*-test, *N* = 47, same participant sample as during initial grid analysis, contrast parametric modulation through entorhinal grid magnitude during observation > baseline, *p* < 0.05 FWE-corrected at cluster level using a cluster-defining threshold of *p* < 0.001, cluster size = 68 voxels; **Figure 4B**). Put differently, striatal activation was stronger when participants observed the demonstrator walking aligned with their individual entorhinal grid orientation. There were no significant activation changes during navigation periods.

We next explored whether such grid-related activation increases were associated with individual variations in performance across participants. In line with our result of increased observation-based grid-like codes in the entorhinal cortex at lower performance (**Figure 3D**), we found significantly increased activation in the bilateral putamen and caudate nucleus (left: *x* = -18, *y* = -2, *z* = 12, *z*-value = 3.72, 222 voxels; right: *x* = 14, *y* = 4, *z* = 18, *z*-value = 3.68, 245 voxels), as well as in the right orbitofrontal cortex (*x* = 32, *y* = 54, *z* = -9, *z*-value = 4.18, 197 voxels) that was time-locked to fluctuations in entorhinal grid magnitude (i.e., stronger when participants observed the demonstrator walking aligned with their individual entorhinal grid orientation) and larger the lower participants performed when re-tracing the demonstrator’s paths (linear regression, *N* = 47, same participant sample as during initial grid analysis, contrast parametric modulation through entorhinal grid magnitude during observation > baseline, cumulative distance error across all paths added as a covariate of interest; note that these results reached significance only at a more lenient statistical threshold, *p* < 0.05 FWE-corrected at cluster level using a cluster-defining threshold of *p* < 0.005, cluster size = 157 voxels; **Figure 4C**). To restate, the activation in these regions scaled more strongly with entorhinal grid-like codes during observation the worse participants performed in re-tracing the demonstrator’s paths thereafter. There were no significant activation changes related to better navigation performance.

### Entorhinal connectivity with striatum and right posterior parietal cortex is time-locked to grid-like codes during observation and negatively scales with performance

So far, we presented evidence that grid-like codes in the entorhinal cortex were significantly enhanced during observation periods and that striatal activation was time-locked to such entorhinal signals. These results were stronger the less accurately participants performed when re-tracing the demonstrator’s paths during navigation periods. To tackle entorhinal-cortical interactions (Gerlei et al., 2021) potentially underlying socio-spatial navigation, we tested functional connectivity changes during observation and navigation. More specifically, we asked whether entorhinal grid-like codes would trigger changes in network connectivity whenever participants would observe the demonstrator walking aligned with their individual grid orientation, and whether such changes would be associated with individual differences in performance.

We tested this using the above model (GLM3) and performed generalized psychophysiological interaction analysis (gPPI, **Materials and Methods**). In brief, we took the anatomical boundaries of the bilateral entorhinal cortex as a seed, calculated its whole-brain connectivity during observation (and navigation) periods (i.e., contrasting the parametric modulation regressor that captured fluctuations time-locked to participant- and path segment-specific entorhinal grid magnitudes with the implicit baseline), and tested whether changes in connectivity varied as a function of the average cumulative distance error per participant.

Results showed that functional connectivity between the entorhinal cortex with the hippocampus and parahippocampal cortex, as well as with the inferior temporal and left lateral prefrontal cortex positively scaled with entorhinal grid magnitudes and that this relationship was stronger at lower performance [linear regression; *N* = 47 (same participant sample as during initial grid analysis), contrast parametric modulation through entorhinal grid magnitude during observation > baseline, cumulative distance error across all paths added as a covariate of interest, *p* < 0.05 FWE-corrected at cluster level using a cluster-defining threshold of *p* < 0.001, cluster size = 61 voxels; **Figure 4D, Table S4**]. This connectivity profile further included enhanced entorhinal coupling with the bilateral striatum and the right posterior parietal cortex (see also **Figure S5** showing the overlap with the right posterior parietal cluster from the grid-independent activation analysis that we reported in **Figure 2C**). Again, these findings appeared specific to lower performance (there were no significant connectivity changes related to better performance) and to observation periods (there were no significant effects during navigation). General connectivity changes (independent of grid-like coding) further showed that entorhinal-cortical coupling was increased during both observation and navigation periods (but stronger during observation; **Figure S6, Table S5**).

Taken together, functional connectivity between the entorhinal cortex, striatum, right posterior parietal cortex and a wide-spread set of cortical regions was time-locked to fluctuations in entorhinal grid magnitudes when participants were tracking the demonstrator’s paths during observation periods and this relationship was stronger at lower individual performance.

## Discussion

In the current study, we investigated whether grid-like codes in the human entorhinal cortex supported socio-spatial navigation as participants tracked and subsequently re-traced the paths of a virtual demonstrator. Our key findings advance current knowledge in several ways: We found that entorhinal grid-like codes supported the tracking of another individual’s movement through space. Crucially, these signals were decoupled from self-movement as participants’ viewpoint remained stationary while observing the demonstrator. Fluctuations in grid magnitudes were associated with the co-activation of striatal regions during observation and these results appeared stronger the less accurate participants performed when subsequently re-tracing the demonstrator’s paths. This profile of co-activation was paralleled by entorhinal connectivity increases with the striatum and the hippocampus, parahippocampal, right posterior parietal and lateral prefrontal cortices when participants observed the demonstrator walking aligned with their internal grid orientations, and this pattern was again more pronounced the worse participants performed. These findings are thus the first to demonstrate that grid-like signaling in the human entorhinal cortex is related to the spatial tracking of others, that this is linked to network dynamics, and modulated by individual task performance.

The main goal of our research was to probe whether grid-like codes in the entorhinal cortex supported the tracking of others, potentially yielding insights into human socio-spatial navigation. Confirming our hypothesis, results showed a significant increase in grid magnitudes (a proxy for putative grid-cell-related activity) while participants were observing the demonstrator’s paths from a stationary viewpoint (**Figure 3B**). This was specific to hexadirectional coding (i.e., a 6-fold, grid-like modulation of the fMRI signal), specific to the entorhinal cortex where grid cells were previously detected (Hafting et al., 2005), and specific to spatial information rather than being driven by eye movements (Julian et al., 2018; Killian et al., 2012; Nau et al., 2018; Staudigl et al., 2018). Several recent findings suggest that socio-spatial navigation relies on neuronal substrates similar to those that process self-related spatial navigation. For instance, Stangl and colleagues (Stangl et al., 2021) used intracranial recordings from the human MTL to reveal boundary-anchored representations during self-navigation as well as during the observation of others moving through space, highlighting a neural mechanism that signals an individual’s vicinity to environmental borders. Further evidence comes from animal work with bats (Omer et al., 2018) and rodents (Danjo et al., 2018) showing so-called “social” place cells in the hippocampus that were specifically tuned to the location of others in space. In contrast, separate subpopulations of CA1 pyramidal cells exclusively coded for self-location or showed a conjoint firing pattern for both self- and other-related location information. Besides hippocampal place cells, grid cell activity (or grid-like coding) is considered central to spatial (Doeller et al., 2010; Hafting et al., 2005; Jacobs et al., 2013; Stangl et al., 2018) and mental navigation (Bellmund et al., 2016; Horner et al., 2016) but was so far not discussed in the context of socio-spatial navigation. Our findings suggest that grid-like codes in the human entorhinal cortex are indeed involved in such a process, allowing us to learn about the spatial routes that others take, and so potentially contributing to our ability to maneuver through crowded and dynamically changing environments as we encounter them in everyday situations.

On the broader scale of cortical structures, flexible self-navigation through the physical environment has been associated with an ensemble of brain regions that includes MTL structures, posterior-medial, parietal, striatal and prefrontal areas (Chersi and Burgess, 2015; Ekstrom et al., 2017; Epstein et al., 2017; Maguire et al., 1998; Patai and Spiers, 2021). Our results are in line with this notion as we found that observing a demonstrator moving through space was associated with increased activation in the hippocampus and adjacent MTL, the caudate nucleus, as well as the ventromedial prefrontal and posterior cingulate cortices (contrast observation > navigation, **Figure 2A**). The hippocampus is regarded as crucial for spatial navigation (Epstein et al., 2017; Maguire et al., 1998, 2000; Morris et al., 1982; O’Keefe and Dostrovsky, 1971), and is assumed to work in parallel with the dorsal striatum (including the caudate nucleus) which was associated with response-learning and goal-directed navigation (Chersi and Burgess, 2015; Doeller et al., 2008; Iaria et al., 2003), as well as route following (Hartley et al., 2003). Especially the latter aspect seems important in the context of the present task since participants were asked to observe, memorize, and to re-trace the paths of a virtual demonstrator akin to following spatial routes. The ventromedial prefrontal and posterior cingulate cortices were previously linked to memory (Bird et al., 2015; King et al., 2015; Rugg and Vilberg, 2012; Wagner et al., 2015, 2016; Watrous et al., 2013), and social cognition (Amodio and Frith, 2006; Gaesser et al., 2019; Gallagher et al., 2000, 2002; Saxe and Powell, 2006; Spreng and Mar, 2012; Wagner et al., 2020), likely supporting memory encoding of the different paths in this socio-spatial setting. Observation additionally triggered activation changes in the posterior superior temporal sulcus and fusiform gyrus, which have both been shown to process biological motion (Herrington et al., 2011; Sokolov et al., 2018). Thus, we revealed a profile of brain regions specifically concerned with tracking others’ complex and goal-directed movements through space.

Evidence on how such macro-scale activation profiles might interact with the putative “spatial map” supported by entorhinal grid codes is sparse. The entorhinal cortex is located at the interface of the hippocampal-neocortical information processing system (Muñoz and Insausti, 2005; van Strien et al., 2009). Its medial entorhinal division, which houses grid cells (Hafting et al., 2005), receives anatomical projections from the hippocampus (van Strien et al., 2009), the parahippocampal (or postrhinal cortex in rodents; Burwell and Amaral, 1998), retrosplenial and posterior parietal cortices (Kerr et al., 2007), as well as from prefrontal areas (Garcia and Buffalo, 2020; Kondo and Witter, 2014). In turn, output from the medial entorhinal cortex is routed back towards the hippocampus (Brun et al., 2008; Cholvin et al., 2021; Remondes and Schuman, 2004) and to a distributed set of cortical areas (Insausti et al., 1997; Muñoz and Insausti, 2005). As such, the entorhinal cortex seems ideally positioned to integrate information and to communicate the current layout of the “spatial map” to the brain-wide navigation network (Gerlei et al., 2021). In the present study, we attempted to characterize whether the activity of entorhinal grid-like codes was time-locked to co-activation of and to communication with regions involved in spatial navigation and visuospatial processing. Results indeed yielded increased co-activation of the striatum (caudate nucleus and putamen) when participants observed the demonstrator walking aligned with their individual entorhinal grid orientations and this effect was stronger the lower participants performed when subsequently re-tracing the paths (**Figure 4BC**). Going further, we next tested for functional interactions and found increased entorhinal connectivity with the caudate nucleus and hippocampus, the parahippocampal, posterior parietal and left lateral prefrontal cortex at larger grid magnitudes during observation, again negatively scaling with performance across participants (**Figure 4D**). Chen and colleagues (Chen et al., 2021) recently investigated the relationship between grid-like codes in the entorhinal and ventromedial prefrontal cortex using human intracranial recordings during virtual, self-related spatial navigation. The authors could show that prefrontal theta oscillations exhibited a hexadirectional signal modulation similar to theta power in the entorhinal cortex and that both types of grid-like codes revealed a comparable grid orientation. These results resonate with our findings of increased co-activation and entorhinal-cortical connectivity as participants observed another individual walking aligned with their internal grid orientation. Altogether, this suggests a mechanism of time-locked network dynamics that are triggered by the activity of grid-like codes in the entorhinal cortex, potentially coordinating information transfer about the current “socio-spatial map” across the brain, informing the observer about how to navigate space either by recalling their own movements, or by learning through observation, as shown here.

Our analyses revealed several surprising findings as well. For instance, we found that entorhinal grid-like codes, as well as their co-activation and entorhinal-cortical connectivity patterns, were positively, rather than negatively, associated with performance (i.e., distance error) in the modified navigation task. In other words, participants with higher grid magnitudes during observation performed worse when re-tracing the demonstrator’s paths (**Figure 3D**). We speculate that accurate performance might go hand-in-hand with neuronally efficient processing (Neubauer and Fink, 2009), which would be indexed by lower entorhinal grid magnitudes and a reduced need to employ additional co-activation of and connectivity with other brain regions. To provide an example, memory training was shown to decrease brain activity during memory processing while increasing performance (Wagner et al., 2021). Individual inability to accurately memorize and re-trace paths might thus be coupled to higher grid-like signals as there was increased need for support through the cognitive map provided by the entorhinal cortex. Notably, previous research reported mixed results, showing increased grid-like codes at better spatial memory performance (measured by a smaller drop-error when trying to place objects at their correct locations; Doeller et al., 2010; Kunz et al., 2015; Stangl et al., 2018), or not showing any significant association (Horner et al., 2016; Nau et al., 2018). Understanding the behavioral implications as well as the exact mechanistic contributions of grid-like codes for observation and navigation thus requires further scrutiny. Moreover, our grid-independent analyses showed that increased activation levels in the right posterior parietal cortex during observation were related to better performance when re-tracing the paths thereafter (**Figure 2C**). The posterior parietal cortex was implicated in converging information about target location, movement, and the position of body parts to be able to plan or make movements towards a target (Whitlock et al., 2008), as well as in tracking spatial routes (Nitz, 2006). This activation cluster was only partly overlapping with grid-modulated voxels (**Figure S5**) but highlights a possible dissociation between overall activation levels and grid-like codes within an anatomical region, and how those signals support performance.

Contrary to previous work (Bellmund et al., 2016; Doeller et al., 2010; Horner et al., 2016; Kunz et al., 2015; Stangl et al., 2018), we did not find significant entorhinal grid-like codes during self-navigation (**Figure 3E**; but see **Figure S3** for significant grid-like coding in the anterior thalamus). The spatial stability of entorhinal grid-like codes was significantly decreased during navigation compared to observation periods (**Figure 3F**), indicating that grid orientations in the entorhinal cortex were less clustered towards a specific direction. Grid orientations hence displayed larger variability across voxels, which might have resulted in an overall decrease in grid magnitude values during navigation. The finding might also be explained by the specific setup of our modified navigation task since participant’s viewpoints were directly placed behind the demonstrator’s starting positions (but note that these randomly varied across trials). This design aspect could have enforced egocentric (striatal) rather than allocentric (hippocampal) processing (Chersi and Burgess, 2015), and could explain the striatal co-activation and connectivity time-locked to observation-related entorhinal grid-like codes. We speculate that decoupling the participant’s viewpoint from the starting position might have rendered the task more hippocampal dependent (Morris et al., 1982). In contrast to this argument, however, it appears that participants formed an allocentric representation since we detected entorhinal grid-like codes during observation, and since results showed generally increased activation levels in the hippocampus and entorhinal cortex during navigation (**Figure S2**). Future research might resolve this issue by expanding our current task-setup and decoupling view- and starting points.

On a final note, we would like to discuss two potential limitations of the present study. First, the design of the modified navigation task did not allow us to disentangle the tracking of others through space from participants planning their own future paths. Prior work demonstrated that entorhinal grid-like codes could be detected as participants imagined movement through space while they remained stationary (Bellmund et al., 2016; Horner et al., 2016). It is possible that participants were imaging their own movements while observing the demonstrator and that this mental navigation caused elevated grid-like codes during observation. However, we consider this an unlikely explanation of the results. Participants completed a post-MRI interview regarding their individual strategies that they had adopted during observation periods. From the 47 participants that were included in the main grid analysis, only one person reported to have imagined the demonstrator’s perspective. We excluded this participant from the sample and repeated the main analysis which left the results unchanged (**Results S5**, and also see this section for a general description of the reported strategies). We acknowledge that this does not fully preclude that processes related to planning and mental navigation contributed to our findings. Future studies might adjust our task design to disentangle potential grid-like processes related to planning, mental navigation, and observation.

Second, while it is plausible that entorhinal grid-like codes support socio-spatial navigation, we would like to emphasize that we are unable to make claims about the social specificity of our results. Entorhinal grid-like codes during observation might also be triggered by non-social features that need to be tracked (such as moving cars when crossing the road) and might be related to feature relevance (keeping an eye on the moving car is important to cross the road safely). While such results would speak for a general role of entorhinal grid-like codes in tracking moving features (Wilming et al., 2018), the debate on the social specificity of brain processes (Lockwood et al., 2020) is fueled by initial evidence for “social” and “non-social” place cells (Omer et al., 2018). Omer and colleagues dissociated signals by either presenting another individual (a demonstrator bat) or an object (a football) moving through space while the observer bat remained stationary. A similar task design might be helpful to resolve this issue in follow-up work with humans. Hence, we cannot draw firm conclusions regarding the involvement of entorhinal grid-like coding specifically in socio-spatial navigation but consider our work an important first step in this direction.

To summarize, we found that grid-like codes in the human entorhinal cortex track other individuals that navigate through space. Grid signals were tied to increases in co-activation and entorhinal-cortical connectivity with an ensemble of regions, including the striatum, hippocampus, parahippocampal and right posterior parietal cortices, altogether modulated by accuracy when subsequently re-tracing the paths. These findings suggest that grid-like codes might be involved in socio-spatial navigation, concerned with tracking others’ complex and goal-directed movements through space. We suggest that grid-like codes and their associated network dynamics could serve to distribute information about the “socio-spatial map” throughout the brain, laying the foundation for an internal “compass” that enables us to maneuver through crowded and dynamically changing (social) environments as we encounter them in everyday situations.

## Acknowledgements

The authors would like to thank Ronald Sladky for support with the MRI sequences and Magdalena Boch for advice on the eye-tracker setup. The project was funded by a grant awarded to I.C.W. [Austrian Science Fund (FWF): P 34775-B].

## Author contributions

Conceptualization: I.C.W. and D.B.O.; Methodology: I.C.W. and M.S.; Software: I.C.W., M.S., and A.L.; Validation: I.C.W. and L.P.G.; Formal Analysis: I.C.W.; Investigation: I.C.W., B.T., and L.P.G.; Resources: I.C.W. and C.L.; Data Curation: I.C.W.; Writing – Original Draft: I.C.W.; Writing – Reviewing & Editing: all authors; Visualization: I.C.W.; Supervision: I.C.W.; Project Administration: I.C.W.; Funding Acquisition: I.C.W.

## Materials and Methods

### Participant sample

Sixty participants volunteered for this study (aged 18-29 years, mean = 22 years, 45 females, 7 right-handed). All participants were healthy, had normal or corrected-to-normal vision, gave written informed consent prior to participation, and received monetary compensation. Two participants were excluded from data analyses (one participant due to an anatomical brain abnormality, and one participant due to low performance in the modified navigation task), which left 58 participants for all following analyses (aged 18-28 years, mean = 22 years, 44 females, 7 right-handed). The study was reviewed and ethically approved by the institutional review board of the University of Vienna (Vienna, Austria, reference number 00538).

### Study procedures and task

Each participant underwent a single MRI session, starting out with the acquisition of the structural brain images while completing the task familiarization period (participants trained the subsequent modified navigation task during one run, all completed the same trials). This was followed by four runs of the actual modified navigation task and the acquisition of functional brain images.

#### Virtual Reality environment and modified navigation task

To investigate whether grid-like codes supported the tracking of others, we translated previous animal work (Omer et al., 2018) to the human context. Our task projected participants into a first-person perspective within a virtual reality (VR) environment and asked them to observe a demonstrator moving through a circular arena. The path had to be held in memory and had to be re-traced after a short delay (for an example video, see https://osf.io/mhtgp/?view_only=ff7038b65c0345cd9ffa4ccd3813d7ba; *note that the OSF project website is currently set to private and will be made publicly available upon publication of the manuscript*).

The VR environment depicted a desert scene with landmarks (sandstone towers, Joshua trees, cacti and other desert plants) and objects (railway wagons, car wrecks, an old gas station, wooden sheds, barrels, water towers, and wind turbines) that were randomly placed around a circular arena. The sandy ground was contrasted by a blue sky filled with clouds, while the sun was fixed at the circular arena’s zenith. The circular arena consisted of a movement area inside which the avatar (i.e., the demonstrator) and the participants could walk (radius = 60 virtual meters, vm; marked in red), as well as an observation area surrounding it (radius = 90 vm, marked in white; **Figure 1A**). All landmarks, objects, and avatars were retrieved from the Unity asset store (https://assetstore.unity.com) and Sketchfab (https://sketchfab.com). Movement speed of both the demonstrator and participant was set to 15 vm/sec and it was not possible to make rotational and translational movements at the same time (it was only possible to walk straight lines but not curves). The rotation speed was set to 50 deg/sec and the participant’s camera height was fixed to 1.7 vm. The VR environment and task were developed using Unity (software version 2019.4.5f1, https://unity.com).

Each trial consisted of an observation and a navigation period (**Figure 1B**). Starting out, a cue signaled participants to pay attention to the demonstrator’s upcoming path (2 seconds, “Please pay attention to the person and copy the path afterwards!”). To be able to visually follow the demonstrator’s path throughout the entire movement area, the observer (i.e., the participant) was projected onto the border of the surrounding observation area, directly placed behind the demonstrator’s starting point. The observer remained stationary during the entire observation period (i.e., the observer could not move or rotate but was able to see the entire movement area) since our goal was to disentangle potential grid-like codes supporting the tracking of others from those associated with self-related spatial navigation. A path consisted of three path segments between successive locations, each segment with a random length between 60-120 vm. Thus, single segments could be walked within 4-8 seconds and an entire observation period could last between 18-30 seconds (consisting of three consecutive segments, the rotation periods, plus a 1.5 s duration before/after the path was started/concluded). This was followed by a jittered delay during which a fixation cross was presented on the computer screen (ranging between 2-7 seconds, mean = 5 seconds).

At the start of the navigation period, the participant (i.e., the former observer) was projected onto the demonstrator’s starting point and was asked to re-trace the path. Using an MR-compatible button box, participants could adjust their orientation and, once confirmed through button press, automatically walked to the opposite border of the movement area (it was not possible to pause and re-adjust inside the movement area). Participants were instructed to follow the previously observed path as close as possible without spending too much time on orientation adjustments. After 30 seconds, a time warning appeared (a red frame was shown around the screen), signaling that participants needed to reach the path’s end point within the next 10 seconds.

Once the border of the movement area was reached for the third time, performance was quantified as (Euclidian) cumulative distance error (vm) averaged across all three points that were visited along the circumference of the movement area (thus, the average difference between the correct and actually visited points). Participants then received feedback about their navigation performance (1 second; happy emoji, distance error =< 20 vm; sad emoji, distance error > 20 vm). This was followed by another jittered delay (fixation cross presented on computer screen, duration ranging between 2-7 seconds, mean = 5 seconds), and a new trial started.

Participants engaged in four task runs of 12 trials each (i.e., consisting of 12 observation and 12 navigation periods). One run had a maximum duration of approx. 17 minutes.

#### Path randomization

Each of the demonstrator’s paths was oriented along one of 36 directions that could be divided into 6 directional bins, spanning the 360°-space with 10° angular resolution (see also **Figure S4**). To estimate individual grid orientations in unbiased fashion (Doeller et al., 2010), we maximized the distance walked (vm) across the different directional bins in two steps.

First, a complete path within one trial consisted of three consecutive path segments. To evenly sample directions from the different directional bins, each individual segment was drawn from a different directional bin (thus, a trial included three path segments sampled from three different directional bins). Across the 12 trials per run (3 path segments per trial), we thus sampled from each directional bin 6 times (6 × 6 = 36 segments per run). Participants completed a total of four runs of the navigation task; each directional bin thus appeared 24 times throughout the experiment. Second, to maximize the distance walked per path segment, we divided the movement area into six equally-sized sectors. Consecutive segments were randomized so that transitions between directly neighboring sectors were prohibited, enforcing a minimum path segment length of 60 vm (the length of a given path segment could vary between 60-120 vm). Starting points of paths were randomly generated within a sector and each sector hosted a start position twice within the same run.

Based on these restrictions, we created two trial sequences that were counter-balanced across the full sample of 60 participants. Overall, there was no significant difference in the distance walked across the 6 directional bins [one-way ANOVA, *N* = 48 (each data point indicating the trial-wise distance walked in each of the directional bins and in each of the four task runs), no significant main effect of directional bin, *p*_two-tailed_ = 0.896, see **Figure S4A**). The specific order of trials was randomly shuffled for each participant.

### Eye tracking acquisition and data processing

To account for the potential impact of eye movements on grid-like codes (Julian et al., 2018; Killian et al., 2012; Nau et al., 2018; Staudigl et al., 2018), and to validate that participants were attending to the demonstrator’s paths, we recorded horizontal and vertical eye gaze, as well as pupil size of each participant’s right eye using a video-based infrared eye tracker (EyeLink 1000 Plus, SR Research, Ontario, Canada). Prior to each recording, raw eye movement data was mapped onto screen coordinates by means of a calibration procedure. Participants sequentially fixated on nine fixation points on the screen, arranged in a 3 × 3 grid. This was followed by a validation procedure during which the nine fixation points were presented once more while the differences between the current and previously obtained gaze fixations (from the calibration period) were measured. If these differences were < 1° of visual angle, the calibration settings were accepted and the eye tracker recording was started. During recording, the data was digitized at a sampling rate of 1000 Hz and a potential drift in eye movements was corrected for every four trials (i.e., approx. every 1.4 minutes). Due to technical problems eye tracking was only possible in a subsample of 47 participants.

Data was corrected for eye blinks by removing samples for which the pupil size deviated more than one standard deviation (s.d.) from the mean across the entire time series. To determine saccadic eye movements, vertical and horizontal eye gaze was transformed into velocities (implemented using Fieldtrip’s “ft_detect_movement”, https://www.fieldtriptoolbox.org). In brief, velocities exceeding a threshold of 6 × the s.d. of the velocity distribution and exceeding a duration of 12 ms were defined as saccades (Engbert and Kliegl, 2003). Saccade onsets during individual path segments (while observing or navigating, excluding any standing or rotation periods) were extracted. To avoid potential artifacts from other eye movements, and since eye movements typically occur every 200-300 ms (Rayner et al., 2009), only events free of saccades and blinks in a 200 ms-interval prior to saccade onset were included (in other words, only events with a pre-saccadic fixation period were considered). We detected a total of 8720 saccades across all participants, across the four task runs, and across both observation and navigation periods (*N* = 47, mean number of saccades ± s.d., overall: 181.7 ± 57.7; observation periods: 97.2 ± 36; navigation periods: 85.45 ± 27.9).

### Imaging parameters

All imaging data were collected at the Neuroimaging Center of the University of Vienna, using a 3T Skyra MR-scanner (Siemens, Erlangen, Germany) equipped with a 32-channel head coil. During each of the four task runs, we acquired on average 474 (± 7, s.d.) T2*-weighted blood oxygenation level-dependent (BOLD) images, using a partial-volume echo-planar imaging (EPI) sequence with the following parameters: repetition time (TR) = 2.029 s, echo time (TE) = 30 ms, number of slices = 30 axial slices, slice order = interleaved acquisition, field of view (FoV) = 216 mm, flip angle = 90°, slice thickness = 3mm, in-place resolution = 2 × 2 mm, using parallel imaging with a GRAPPA acceleration factor of 2. Slices were oriented parallel to the long axis of the hippocampus.

Since the entorhinal cortices are susceptible to image distortions due to their vicinity to air-filled cavities, we acquired 30 images for post-hoc artifact correction using the abovementioned functional sequence but reversing the phase-encoding direction (thereby stretching potential image distortions into the opposite direction). Additionally, to facilitate the co-registration of anatomical entorhinal cortex masks to the partial-volume EPI images, we acquired 10 whole-brain EPI images with the following parameters: repetition time (TR) = 2.832 s, echo time (TE) = 30 ms, number of slices = 42 axial slices, slice order = interleaved acquisition, field of view (FoV) = 216 mm, flip angle = 90°, slice thickness = 3mm, in-place resolution = 2 × 2 mm, using parallel imaging with a GRAPPA acceleration factor of 2. This sequence was thus similar to the partial-volume EPIs (but had a longer TR to allow for whole-brain coverage). As above, slices were oriented parallel to the long axis of the hippocampus.

The T1-weighted structural image was acquired using a Magnetization-Prepared Rapid Gradient Echo (MPRAGE) sequence with the following parameters: TR = 2.3 s; TE = 2.43 ms; FoV = 240 mm, flip angle = 8°, voxel size = 0.8 mm isotropic. To delineate the entorhinal cortex, we acquired a T2-weighted structural image using a turbo-spin-echo (TSE) Sampling Perfection with Application optimized Contrasts using different flip angle Evolution (SPACE) sequence with the following parameters: TR = 3.2 s; TE = 564 ms; FoV = 256 mm, voxel size = 0.8 mm isotropic. These slices were oriented perpendicular to the long axis of the hippocampus.

### MRI data preprocessing

The MRI data were processed using SPM (software version 12, http://www.fil.ion.ucl.ac.uk/spm/) in combination with Matlab (software version R2019b, The Mathworks, Natick, MA, USA) and the Functional Magnetic Resonance Imaging of the Brain (FMRIB) Software Library (FSL, software version 5.0.1; https://fsl.fmrib.ox.ac.uk/fsl/fslwiki/; Jenkinson et al., 2012). The first six volumes were excluded to allow for T1-equilibration. The remaining volumes were slice-time-corrected to the middle slice and realigned to the mean image calculated across the four task runs. Potential image distortions were corrected by applying FSL’s “topup*”* command: we calculated the mean image based on the additional volumes acquired (phase-encoding direction reversed). Together with the original fMRI data, this image was then used to estimate and correct susceptibility-induced distortions. Since grid-like codes were analyzed in the participant-specific image space, we refrained from normalizing the data but applied a 3D Gaussian smoothing kernel (5 mm full-width at half maximum, FWHM).

For the whole-brain group analyses, the distortion-corrected data was additionally normalized into standard space. The structural scan was co-registered to the mean functional image and segmented into grey matter, white matter, and cerebrospinal fluid using the “New Segmentation” algorithm. All images (functional and structural) were spatially normalized to the Montreal Neurological Institute (MNI) EPI template (MNI-152) using Diffeomorphic Anatomical Registration Through Exponentiated Lie Algebra (DARTEL, Ashburner, 2007), and functional images were smoothed with a 3D Gaussian kernel (5 mm FWHM).

### Whole-brain univariate fMRI analysis

We first set out to test whole-brain activation changes during observation and navigation, independent of grid-like codes. Using a Generalized Linear Modeling (GLM) approach, the BOLD response during the modified navigation task was modeled using separate task regressors, time-locked to the onsets of the respective events (cues, observation periods, navigation periods, feedback). All events were estimated as boxcar functions of specific durations and were convolved with the SPM default canonical hemodynamic response function (HRF). Cue and Feedback periods were modelled with a duration of 2 and 1 seconds, respectively. The duration of observation and navigation periods varied depending on the path length and the participant’s behavior, and was defined through the on- and offsets of the VR environment on the computer screen (ranging between 18-40 seconds; see above). This included events such as orientation adjustments (rotations), walked path segments (translation periods), and time periods during which no movement occurred (short standing periods in-between). To account for noise due to head movement, we included the six realignment parameters, their first derivatives, and the squared first derivatives into the design matrix. A high-pass filter with a cutoff at 128 s was applied. The four runs of the modified navigation task were combined into one first-level model and contrasts were created ([observation ∩ navigation] > implicit baseline, observation/navigation > implicit baseline, observation > navigation and vice versa), collapsing across the different runs. To test for group effects, these contrast images were submitted to one-sample *t*-tests.

Additionally, we were interested in whether activation changes during observation and/or navigation scaled with individual performance. We thus ran two linear regression analyses (contrasts observation/navigation > implicit baseline) and added the cumulative distance error (vm, obtained during navigation periods and averaged across all three points of a path trajectory) as a covariate of interest.

### Definition of regions-of-interest (ROIs)

#### Entorhinal cortex ROIs and participant exclusions

We used the T2-weighted structural scans to anatomically delineate individual entorhinal cortices. First, ROIs were automatically generated using Automated Segmentation of the Hippocampal Subfields (ASHS, software version 1.0.0, https://sites.google.com/site/hipposubfields/; Yushkevich et al., 2015). Second, to verify the ASHS-based segmentation, we also performed manual delineation of the entorhinal cortex by tracing its anatomical borders on the structural image. This was done using ITK-SNAP (software version 3.6.; www.itksnap.org; Yushkevich et al., 2006), following the segmentation protocol provided by Berron and colleagues (Berron et al., 2017).

As we initially did not have a specific hypothesis regarding the laterality of brain effects, we collapsed the left and right masks into a bilateral entorhinal cortex image (for both the ASHS- and the ITK-SNAP-based delineations; but see **Results S2** and **S3** for separate analyses of ROIs in the left and right hemispheres). These masks were then binarized and transformed into the participant-specific space of the functional images. Since the functional images were only partial-volume slabs, co-registration was aided by an additional intermediate step that involved the mean whole-brain functional image (Stangl et al., 2018). First, each participant’s T2-weighted structural image (together with the individual entorhinal mask) was co-registered to match the orientation of the mean whole-brain functional image. Second, the mean whole-brain functional image (together with the co-registered individual entorhinal mask) was co-registered to match the orientation of the mean partial-volume functional image (mean ± s.d.; ASHS, 56 ± 13 voxels; ITK-SNAP, 104 ± 23 voxels).

The quality of co-registration was confirmed through visual inspection of each mask’s overlap with the individual (co-registered) structural and functional data. The entorhinal cortex lies in close proximity to the temporal horn of the lateral ventricle. Such tissue borders are often associated with lower signal-to-noise ratio, which is also what we experienced in a subsample of participants. To circumvent this issue, we only considered voxels that exceeded a signal-to-noise threshold of 0.8, leading to the fact that voxels along the anterior-medial entorhinal cortex border were partly dropped from the analyses (participants were excluded if there were less than 5 voxels left in the mask, and two participants were fully excluded from all grid code analyses involving the entorhinal cortex). After applying these restrictions, the final participant sample for which entorhinal cortex data was available comprised 49 (ASHS; 23 ± 13 voxels) or 51 (ITK-SNAP; 38 ± 25 voxels) participants.

#### Control ROIs

To test whether grid-like codes were also detectable in other regions, we chose several control ROIs known to be involved in spatial navigation and visual processing but for which no significant grid-like coding was reported so far. These included the adjacent hippocampus, the parahippocampal cortex, the anterior thalamus, and the primary visual cortex (V1). Both the hippocampus and parahippocampal cortex masks were defined using the ASHS algorithm (the hippocampus was defined by merging the hippocampal subfields cornu ammonis (CA) 1-4 and the subiculum). To delineate the anterior thalamus, we used the stereotactic mean anatomical atlas provided by Krauth and colleagues (Krauth et al., 2010)(© University of Zurich and ETH Zurich, Axel Krauth, Rémi Blanc, Alejandra Poveda, Daniel Jeanmonod, Anne Morel, Gábor Székely), which is based on histological, cytoarchitectural features defined *ex vivo* (Morel, 2007). We specified the anterior thalamus by combining the anterior dorsal, -medial, and -ventral nucleus masks. The V1 mask was created using the Automatic Anatomical Labeling (AAL) atlas (Tzourio-Mazoyer et al., 2002).

As above, left and right masks were combined into bilateral volumes and were transformed into the participant-specific image space (hippocampus, 447 ± 56 voxels; parahippocampal cortex, 160 ± 26 voxels; anterior thalamus, 41 ± 6 voxels; V1, 1136 ± 297 voxels). The quality of co-registration was confirmed through visual inspection of each mask’s overlap with the individual structural and functional data of each participant (final sample, *N* = 58 participants).

### Analysis of grid-like codes

We next asked whether grid-like codes supported the tracking of others. All analyses were based on the openly available source code of the Grid Code Analysis Toolbox (GridCAT, software version 1.0.4, https://www.nitrc.org/projects/gridcat; Stangl et al., 2017), which follows the procedures established by Doeller and colleagues (Doeller et al., 2010).

#### Estimating grid orientations (GLM1)

Data was modelled identical to above with the exception that each path segment was included as a separate event and was modulated by its direction. In brief, the BOLD response during the modified navigation task was modeled using separate task regressors, time-locked to the onsets of the respective events (cues, path segments observed during observation periods, path segments walked during navigation periods, feedback). Translational events (i.e., individual path segments observed by the participant/walked by the demonstrator, and path segments walked by the participant) were modelled from the start of the movement until the next point was reached (thus, the duration was depended on the path segment length). To obtain the direction of each segment (i.e., translational event *t*), we calculated the translation angle (α_t_) based on the path segment coordinates within the movement area and referenced them to an arbitrary zero coordinate on the horizontal VR plane. The translational direction was then modelled using two parametric modulators, defined as sin(α_t_*6) and cos(α_t_*6). Orientation adjustments (rotations) and time periods during which no movement occurred (standing periods) were not explicitly modelled as these durations were typically very short (< 2 seconds). Grid code analysis was performed in the native-space of each participant.

We then estimated individual grid orientations by partitioning the fMRI data into estimation and test sets. To allow for stable estimations, we maximized the data available for grid orientation estimation by leveraging the inherent trial-design of each run. Specifically, we used a 12-fold cross-validation (CV) regime during which grid orientations were estimated on the path segments of 11 trials (consisting of 11 observation and 11 navigation periods) and tested on the path segments of the remaining trial (consisting of 1 observation and 1 navigation period). This was iterated until every trial was tested once.

During each CV-fold, voxel-wise grid orientations for observation/navigation conditions were estimated by fitting the fMRI data using GLM1. We then calculated the mean grid orientation (ϕ) of each participant based on the average beta estimates (β_1_ and β_2_) of the two parametric modulators, extracted from all voxels within the respective ROI and calculated using arctan[mean(β_1_)/mean(β_2_)]/6.

#### Testing grid orientations (GLM2)

This model was identical to GLM1 but instead modelled each translational event modulated by the difference between the event’s translational direction (α_t_) and the participant’s mean grid orientation (ϕ) using cos[6*(α_t-_ ϕ)]. This analysis yielded a voxel-wise grid magnitude value for each path segment (i.e., the strength of grid-like signal per path segment that was observed or walked).

Finally, results were averaged across CV-folds and across voxels within the respective ROI, and data was analyzed using a set of one-sample *t*-tests. We *a priori* expected that the magnitude of entorhinal grid-like codes should be greater than zero, which is why we adopted an α-level of 0.05 (one-tailed). Additionally, we applied Bonferroni-correction to account for multiple comparisons (2 entorhinal cortex ROIs and 4 control ROIs), using a threshold of α_Bonferroni_ = 0.05/6 ROIs = 0.008. Grid magnitude values exceeding the median value ± 3 × the median absolute deviation were excluded from the analyses. Results remained stable when basing calculations on the full data set (i.e., not excluding any outliers, see **Tables S2** and **S3**) and the raw data is publicly provided (see below).

#### Representational stability: spatial and temporal stability of grid orientations

Grid-like codes might be affected by variations in spatial and temporal signal stability (Stangl et al., 2018). In the case of high spatial instability, the estimated grid orientations might differ across the voxels within a ROI, resulting in random mean grid orientations and decreased grid magnitudes of grid-like codes. In the presence of low temporal stability, decreased grid magnitudes could be caused by varying grid orientations over time (for example, different grid orientations present in the estimation vs. test set).

To estimate the extent to which such stability aspects affected our results, we repeated the abovementioned analysis (GLM1, which served to quantify voxel-wise grid orientations) but partitioned each run into data halves (this way, we were able to estimate temporal stability in equally-sized data parts which would not have been possible with the CV-regime). To obtain a metric of spatial stability, we then calculated the coherence of estimated voxel-wise grid orientations between all voxels within the ROI using Rayleigh’s test for non-uniformity of circular data. Higher *z*-values were associated with higher spatial stability (thus, significantly clustered grid orientations within the ROI). To obtain a metric of temporal stability, we compared the voxel-wise grid orientation between the first and second data half. A voxel was classified as “stable” if the differences in grid orientations was within ± 15° and temporal stability was quantified by the proportion (%) of ‘stable’ voxels within the ROI.

### Brain activation and connectivity time-locked to grid-like codes

Next, we asked whether the activity of entorhinal grid-like codes was time-locked to voxel-wise changes in cortical activation and entorhinal-cortical connectivity, and whether such dynamics were modulated by individual performance. To obtain a value of grid-like codes for each path segment observed or walked, we leveraged GLM2 (which was estimated in each of the 12 CV-folds) and extracted the grid magnitudes from the parametric modulation regressor for each path segment (i.e., relying on the difference between each event’s translational direction and the mean grid orientation of the participant, whereby a smaller difference should be associated with a stronger grid-like signal within the entorhinal cortex ROI that was automatically segmented with ASHS). Data was then modelled similar to above (GLM1, GLM2) but translational events were modulated by the path segment-specific grid magnitudes. To perform group-level analyses, the GLM3 was estimated based on fMRI data in MNI standard-space.

We contrasted the parametric modulation regressors that the captured path segment-wise fluctuations in entorhinal grid magnitude against baseline (entorhinal grid magnitude during observation/navigation > implicit baseline) and tested for group effects by submitting the individual contrast images to separate one-sample *t*-tests. Additionally, to test whether activation changes time-locked to fluctuations in entorhinal grid magnitude scaled with performance, we ran linear regression analyses and added the cumulative distance error (vm, obtained during navigation periods and averaged across all three points of a path) as a covariate of interest.

#### Generalized Psychophysiological Interaction (gPPI) analysis

Moreover, our goal was to probe whether fluctuations in entorhinal grid magnitude (i.e., the strength of the grid-like signal during each path segment) would be associated with entorhinal-cortical functional connectivity changes. To achieve this, we used the abovementioned model (GLM3) and performed generalized psychophysiological interaction analysis (gPPI; McLaren et al., 2012). We took the anatomical boundaries of the bilateral posterior-medial entorhinal cortex as a seed (Maass et al., 2015), extracted the first eigenvariate of its functional timecourse and adjusted for average activation levels using an *F*-contrast. The timecourse was then deconvolved to estimate the putative neural activity of the seed region (i.e., the physiological factor) and was multiplied with boxcar functions that defined the specific task events (i.e., the psychological factor). The resulting vectors were convolved with the canonical HRF, yielding one psychophysiological interaction regressor per condition-of-interest (i.e., for parametric modulation regressors that captured fluctuations in entorhinal grid magnitude during path segments of observation/navigation periods), and were contrasted against the implicit baseline.

Group-level connectivity analyses were performed using a set of one-sample *t*-tests. Again, linear regression analysis was used to test for connectivity changes time-locked to fluctuations in entorhinal grid magnitude that potentially scaled with performance by adding the cumulative distance error as a covariate of interest.

### Statistical analyses and thresholding

#### Statistical analysis of behavioral data and ROI-based fMRI results

Analyses of navigation performance (distance error, vm) and ROI-based fMRI results (magnitude of grid-like codes, directional bins and representational stability) were carried out using R (https://www.r-project.org), using a set of *t*-tests and ANOVA models. Significant interaction effects were followed-up with pair-wise comparisons using the R-package *emmeans* (https://cran.r-project.org/web/packages/emmeans/index.html) and were corrected for multiple comparisons (Tukey’s HSD). Effect sizes were calculated as Cohen’s *d* or eta squared (*η*^*2*^) for *t*-test and ANOVA models, respectively (Lakens, 2013). As mentioned above, we *a priori* expected the magnitude of entorhinal grid-like codes should be greater than zero, which is why we adapted an α-level of 0.05 (one-tailed). Additionally, we applied Bonferroni-correction to account for multiple comparisons, using a threshold of α_Bonferroni_ = 0.05/6 ROIs = 0.008. For all other cases, the α-level was set to 0.05 (two-tailed). Any exploratory analyses are explicitly described as such. Grid magnitude values exceeding the median value ± 3 × the median absolute deviation were excluded from the analyses. Results remained stable when basing calculations on the full data set (i.e., not excluding any outliers) and are reported in **Tables S2** and **S3**.

#### Statistical thresholding of whole-brain fMRI results and anatomical labeling

Unless stated otherwise, significance for all whole-brain fMRI analyses was assessed using cluster-inference with a cluster-defining threshold of *p* < 0.001 and a cluster-probability of *p* < 0.05 family-wise error (FWE) corrected for multiple comparisons. The corrected cluster size (i.e., the spatial extent of a cluster that is required in order to be labeled as significant) was calculated using the SPM extension “CorrClusTh.m” and the Newton-Raphson search method (script provided by Thomas Nichols, University of Warwick, United Kingdom, and Marko Wilke, University of Tübingen, Germany; http://www2.warwick.ac.uk/fac/sci/statistics/staff/academic-research/nichols/scripts/spm/).

Anatomical nomenclature for all tables was obtained from the Laboratory for Neuro Imaging (LONI) Brain Atlas (LBPA40, http://www.loni.usc.edu/atlases/).

## Data availability

All anonymized data are available upon reasonable request to the authors in accordance with the requirements of the institute, the funding body, and the institutional ethics board. Raw data (behavioral performance and ROI-based results of all grid analyses), as well as unthresholded statistical whole-brain fMRI maps are available at the Open Science Framework (https://osf.io/mhtgp/?view_only=ff7038b65c0345cd9ffa4ccd3813d7ba; *note that the OSF project website is currently set to private and will be made publicly available upon publication of the manuscript*).

## Code availability

All analysis is based on openly available software. Software codes for the modified navigation task are available upon reasonable request to the authors.

## Supplementary Materials

### General note on data and materials available on the Open Science Framework

This article is linked to the following data on the Open Science Framework (OSF; https://osf.io/mhtgp/?view_only=ff7038b65c0345cd9ffa4ccd3813d7ba; *note that the OSF project website is currently set to private and will be made publicly available upon publication of the manuscript*): (1) raw behavioral data from the modified navigation task; (2) results from the grid code analysis; (3) brain activation/connectivity maps (both unthresholded and thresholded); (4) corresponding tables with MNI-coordinates (note that, unless stated otherwise, tables included in this supplement only show the peak coordinates of each cluster).

## Supplementary Results

### Results S1: Increased activation in the right posterior parietal cortex during observation is associated with better performance

Since participants generally performed best when re-tracing the first path segment of a given trajectory (**Figure S1**), we reasoned that good performance might rather be reflected by a smaller distance error on the second and third path segments (i.e., the cumulative distance error_S2+S3_). We thus repeated our analysis, regressing the cumulative distance error_S2+S3_ against voxel-wise brain activity. Results were virtually identical, again showing increased activation in the right posterior parietal cortex (*x* = 44, *y* = -78, *z* = 24, *z*-value = 4.5, 226 voxels) during observation that negatively scaled with the individual cumulative distance error across path segments 2-3 [linear regression, *N* = 58, contrast observation > baseline, cumulative distance error_S2+S3_ added as a covariate of interest; *p* < 0.05 family-wise-error (FWE) corrected at cluster level using a cluster-defining threshold of *p* < 0.001, cluster size = 80 voxels].

### Results S2: Entorhinal grid-like codes when observing the demonstrator’s paths

Our main results of significant grid-like codes during observation remained stable when we delineated the entorhinal cortex following a manual segmentation protocol (**Materials and Methods**). In brief, we found significant grid-like codes in the entorhinal cortex as participants were observing the demonstrator’s paths. This effect was only present for the 6-fold symmetrical model [one-sample *t*-test, *N* = 51 (full data set, no exclusions); *t*(50) = 1.9, Cohen’s *d* = 0.27, 95% confidence interval (CI) = [-0.01, 0.18], *p*_one-tailed_ = 0.032, although not surviving correction for multiple comparisons; **Figure S3A;** results for this and all following analyses of this section remained stable when using the full data set, see also **Table S2**]. Different symmetrical models such as 5- or 7-fold signal periodicities did not yield significant results [5-fold: *N* = 47 (excluding 4 outliers), *p*_one-tailed_ = 0.102; 7-fold: *N* = 46 (excluding 5 outliers), *p*_one-tailed_ = 0.178; **Figure S3A**].

#### Significant entorhinal grid-like codes during observation when using a different cross-validation regime

Our initial cross-validation regime involved estimating the putative grid orientation on 11 of 12 trials (GLM1) and testing the signal modulation on the remaining three path segments of the held-out trial (GLM2). To assure that the parametric modulation of the held-out data could be reliably estimated, we repeated the main analysis and used a modified, less nested cross-validation regime by splitting each task run into four data parts [i.e. estimating grid orientations on three fourths (27 path segments) and testing on the remaining (9 path segments)]. Results remained stable, showing significantly increased grid-like codes in the entorhinal cortex (defined through automated segmentation using ASHS, **Materials and Methods**) during observation [6-fold symmetrical model, mean grid magnitude (arbitrary units, a.u.) ± s.e.m., 0.077 ± 0.04, one-sample *t*-test, *N* = 45 (excluding 4 outliers); *t*(44) = 2.013, Cohen’s *d* = 0.3, 95% CI = [0, 0.15], *p*_one-tailed_ = 0.025].

#### Grid-like codes during observation appear left lateralized

Since the grid-like modulation of the entorhinal signal was reported to be right-lateralized in previous navigation studies using virtual reality (Doeller et al., 2010), we repeated our analyses but separately tested for entorhinal grid-like codes in the left and right hemisphere (ROIs defined through automated segmentation using ASHS). We used the cross-validation scheme reported above (data was partitioned into four parts for estimating and testing individual grid orientations). Participants with less than 5 voxels within the ROI mask were excluded from the analyses which yielded a final sample of 38 and 37 participants for the left (mean ± s.d., 13 ± 7 voxels) and right (14 ± 7 voxels) hemisphere, respectively.

Findings revealed significant grid-like codes in the left [6-fold symmetrical model, mean grid magnitude (arbitrary units, a.u.) ± s.e.m., 0.078 ± 0.03, one-sample *t*-test, *N* = 37 (excluding 1 outlier); *t*(36) = 2.421, Cohen’s *d* = 0.4, 95% CI = [0.01, 0.14], *p*_one-tailed_ = 0.021] but not in the right entorhinal cortex during observation [6-fold symmetrical model, mean grid magnitude (arbitrary units, a.u.) ± s.e.m., -0.023 ± 0.06, one-sample *t*-test, *N* = 37 (full sample); *p*_one-tailed_ = 0.339].

#### Negative grid-like codes in V1

We further investigated the result of significantly negative grid-like codes in primary visual cortex (V1) during observation. One reason could be the small number of path segments in the test set using our initial 12-fold cross-validation regime (grid orientations were iteratively estimated on 11 trials/33 path segments and tested on 1 trial/3 path segments), whereby observing/moving in single directions could have driven the result (e.g., a strong response in a specific direction compared to smaller responses in other directions). We thus repeated our analysis with a different cross-validation regime (reported above, splitting the data into four parts for estimating/testing grid orientations). However, results remained unchanged and once more revealed significantly negative grid-like coding in V1 during observation [6-fold symmetrical model, mean grid magnitude (arbitrary units, a.u.) ± s.e.m., -0.82 ± 0.11, one-sample *t*-test, *N* = 55 (excluding 1 outlier); *t*(54) = -7.25, Cohen’s *d* = -0.98, 95% CI = [-1.04, -0.59], *p*_one-tailed_ < 0.0001].

Negative grid-like codes (in the entorhinal cortex) were reported previously (Horner et al., 2016; Nau et al., 2018). Considering our cross-validation approach, a significantly negative 6-fold symmetrical modulation of the fMRI signal in V1 could indicate increased grid magnitudes for directions that are rotated 30°, resulting in increased grid magnitudes opposite to the estimated grid orientation. Following an interpretation by Horner and colleagues, such an effect could be due to reduced neuronal firing when participants track others walking aligned with their internal (V1) grid orientation, given that inhibitory mechanisms appear vital for grid-like coding (Couey et al., 2013). At present, however, we do not know which factors drive this effect and cannot provide a firm interpretation of why negative grid-like coding in V1 appears associated with tracking others navigating through space.

### Results S3: No significant entorhinal grid-like codes when re-tracing the demonstrator’s paths

We did not find significant grid-like codes in the entorhinal cortex, also not when manually delineating the region [*N* = 45 (excluding 6 outliers), *p*_one-tailed_ = 0.323; **Figure S3B**; results for this and all following analyses of this section remained stable when using the full data set, **Table S3**], or when testing for grid-like codes in the hippocampus [*N* = 52 (excluding 6 outliers), *p*_one-tailed_ = 0.34], the parahippocampal cortex [*N* = 56 (excluding 2 outliers), *p*_one-tailed_ = 0.087], or V1 [*N* = 52 (excluding 6 outliers), *p*_one-tailed_ = 0.222; **Figure S3B**]. However, we found significantly increased grid magnitudes in the anterior thalamus which was present when applying a 6-fold symmetrical model to the data [*N* = 54 (excluding 4 outliers), *t*(53) = 2.186, *d* = 0.3, 95% CI = [0.02, 0.48], *p*_one-tailed_ = 0.016, although not surviving correction for multiple comparisons; **Figure S3B**].

#### No significant grid-like codes during navigation when using a reduced data set

Additionally, we speculated that the lack of grid-like codes when re-tracing the demonstrator’s paths might have been caused by the specifics our task design. Participants were placed directly behind the demonstrator’s starting position during observation (note that the starting position randomly varied from trial-to-trial). This might have prompted participants to adopt an ego-rather than allocentric reference frame, especially during the first path segment (the remaining path segments varied randomly relative to the participant’s initial viewpoint). If this was the case, we would have expected decreased grid-like codes in the entorhinal cortex when traversing path segments that were decoupled from the initial viewpoint (i.e., path segments 2-3). We thus repeated the above analysis and tested for grid-like codes during navigation when re-tracing the initial vs. the remaining path segments (i.e., collapsing across path segments 2-3; **Materials and Methods**). Results did not show significant grid-like codes in the entorhinal cortex when participants re-traced the initial path segment [one-sample *t*-test; *N* = 42 (excluding 7 outliers), *p*_one-tailed_ = 0.09; *N* = 49 (full sample), *p*_one-tailed_ = 0.156], and also not when they re-traced segments 2-3 [one-sample *t*-test; *N* = 38 (excluding 11 outliers), *p*_one-tailed_ = 0.125; *N* = 49 (full sample), *p*_one-tailed_ = 0.156], confirming our initial findings (however, please note that these results are based on a reduced data set which might have contributed to the lack of significant findings).

#### No significant grid-like codes during navigation when using a different cross-validation regime

As mentioned previously, our initial cross-validation regime involved estimating the putative grid orientation on 11 of 12 trials (GLM1) and testing the signal modulation on the remaining three path segments of the held-out trial (GLM2). Such heavily nested cross-validation might negatively affect the reliable estimation of the parametric modulation in the held-out data set. We thus repeated the main analysis and used a modified, less nested cross-validation regime by splitting each task run into four data parts (see above). Results remained stable, showing no significant grid-like codes in the entorhinal cortex (defined through automated segmentation using ASHS, **Materials and Methods**) during navigation [6-fold symmetrical model, one-sample *t*-test, *N* = 46 (excluding 3 outliers), *p*_one-tailed_ = 0.247].

#### Grid-like codes during navigation appear right lateralized

Following our approach above, we repeated the analyses but separately tested for entorhinal grid-like codes in the left and right hemisphere (ROIs defined through automated segmentation using ASHS; N = 38; mean ± s.d.; left ROI: 13 ± 7 voxels, right ROI: 14 ± 7 voxels; 4-fold cross-validation regime). We found significantly negative grid-like codes in the left [6-fold symmetrical model, mean grid magnitude (arbitrary units, a.u.) ± s.e.m., -0.071 ± 0.04, one-sample *t*-test, *N* = 37 (excluding 1 outlier); *t*(36) = -1.817, Cohen’s *d* = -0.29, 95% CI = [-0.15, 0.01], *p*_one-tailed_ = 0.039] and numerically increased (but not significant) grid-like codes in the right entorhinal cortex during navigation [6-fold symmetrical model, mean grid magnitude (arbitrary units, a.u.) ± s.e.m., 0.085 ± 0.05, one-sample *t*-test, *N* = 37 (excluding 1 outlier); *p*_one-tailed_ = 0.06].

#### Grid orientations during observation did not match those during navigation

We further explored whether grid orientations during observation periods served as spatial reference frames when participants re-traced the demonstrator’s paths during navigation. If this was the case, we would expect to find matching grid orientations between the two conditions, leading to significantly increased grid magnitudes when testing grid orientations obtained from observation- on navigation periods. We repeated the above analysis but estimated individual grid orientations based on all paths during the 12 observation periods of each task run, testing them on all paths during the 12 navigation periods of each task run. Results did not yield significant grid magnitudes in the entorhinal cortex when testing grid orientations obtained from observation- on navigation periods [one-sample *t*-test, *N* = 45 (excluding 4 outliers), *p*_one-tailed_ = 0.206; *N* = 49 (full sample), *p*_one-tailed_ = 0.156; **Figure S3C**] or vice versa [i.e., grid orientations obtained from navigation periods did not match those during observation; *N* = 45 (excluding 4 outliers), *p*_one-tailed_ = 0.46; *N* = 49 (full sample), *p*_one-tailed_ = 0.494; **Figure S3C**].

### Results S4: No effect of eye movements on grid-like codes

We repeated our initial grid analysis but instead modelled eye gaze directions (i.e., the angle between successive saccades with respect to an arbitrary reference point on the computer screen) instead of movement trajectories in VR-space (note that eye-tracking data was available only for a subset of 47 participants; of these, grid code analyses were possible for *N* = 37 participants that had a sufficient amount of voxels in the entorhinal cortex mask; **Materials and Methods**). Each task run was partitioned into data halves to independently estimate and test grid orientations (i.e., individual grid orientations were estimated on all saccades during trials 1-6 and were tested on all saccades during trials 7-12). As above, grid orientations were assessed separately for observation and navigation periods.

Results showed no significant increase in entorhinal grid magnitudes when testing for a 6-fold symmetrical model during observation periods [separate one-sample *t*-tests; *N* = 33 (excluding 4 outliers), *p*_one-tailed_ = 0.433; *N* = 37 (full sample), *p*_one-tailed_ = 0.449]. However, the distance viewed across the different directional bins appeared biased [i.e., participants made more or longer saccades along directional bins 1 and 6 which covered horizontal eye movements between the left/right borders of the observed movement area; repeated-measures ANOVA, *N* = 33, significant main effect of directional bin, *F*(5,186) = 30.1, *η*^*2*^= 0.43, 95% CI = [245.9, 266.5], *p*_two-tailed_ < 0.0001; **Figure S4C**]. Since this could have distorted the estimation of grid orientations, we repeated the above analysis and randomly selected saccades to match the average, individual distance viewed along directional bins 2 to 5. Again, results showed no significant saccade-related grid-like codes in the entorhinal cortex during observation [*N* = 37 (full sample, no outliers), *p*_one-tailed_ = 0.35; **Figure S4C**], and also not during navigation periods [*N* = 35 (excluding two outliers), *p*_one-tailed_ = 0.096; *N* = 37 (full sample), *p*_one-tailed_ = 0.027; **Figure S4C**]. This suggests that our finding of significant grid-like codes in the entorhinal cortex when observing the demonstrator’s paths (**Figure 3B**) was based on spatial rather than visual information.

#### Participants followed the demonstrator with their eye gaze during observation

To verify that participants actually followed the demonstrator with their eye gaze during observation, we demarcated rectangular areas-of-interest (AOIs) that were defined by the two-dimensional coordinates of a given path segment on the computer screen (i.e., its starting and end points) as well as by the demonstrator’s height on the screen (thus, defining the area on the screen within which the demonstrator was moving). We then calculated the percentage of eye movements that fell within the AOI boundaries. We reasoned that participants would produce slower (smooth pursuit) as well as faster (saccadic) eye movements to track the demonstrator and hence based calculations on the raw data that was cleaned from eye blinks (i.e., data was not restricted to saccades only, as was the case for the abovementioned saccade-based grid code analysis).

Indeed, the majority of eye movements were located within AOIs, emphasizing that participants’ viewing behavior was related to the observation of the demonstrator [*N* = 47 (full sample for which eye tracking data was available), percentage of eye movements detected within AOI per path segment, mean ± s.e.m., first path segment: 80.1 ± 2.41 %, second path segment: 77.11 ± 1.9 %, third path segment: 73.21 ± 1.68 %]. There was no difference in the amount of AOI-related eye movements across the three path segments [one-way ANOVA, *N* = 47, no significant main effect of path segment, *p*_two-tailed_ = 0.057].

### Results S5: Reported strategy use during the modified navigation task

After completing the MRI experiment, participants were interviewed regarding their individual strategies that they had adopted during observation periods (“Briefly describe in your own words how you oriented yourself within the virtual reality environment during observation/navigation.”). Of all 58 participants, 16 (27.6%) reported that they payed attention to the landmarks while tracking the avatar and during self-navigation (strategy “landmark”), 30 (51.7%) reported that they payed attention to the landmarks as well as to the magnitude of the movement angles in-between subsequent path segments (strategy “landmark + angle”), 6 (10.3%) reported that they payed attention to the landmarks as well as to the duration of the rotation when the avatar was adjusting its orientation in-between path segments (strategy “landmark + rotation”), 4 (6.9%) reported that they payed attention to the landmarks and tried to imagine themselves in the demonstrator’s perspective during observation (strategy “landmark + demonstrator perspective”), and 2 (3.4%) reported creating a “mental map”, trying to capture the general layout of the entire environment as good as possible (strategy “mental map”)^1^.

It was previously shown that entorhinal grid-like codes could be detected as participants imagined movement through space while they remained stationary (Bellmund et al., 2016; Horner et al., 2016). It is possible that participants were imaging their own movements while observing the demonstrator and that this mental navigation caused elevated grid-like codes during observation. From the 47 participants that were included in the main grid analysis, only one person reported to have imagined the demonstrator’s perspective (the remaining three participants were excluded from this analysis because of outlier values or because of insufficient data within the entorhinal cortex ROI). We excluded this participant from the sample and repeated the main analysis (see **Results** and **Materials and Methods** section; 12-fold cross validation to detect grid-like codes during observation periods, automatic segmentation of the entorhinal cortex using ASHS). Results remained unchanged and once more confirmed our finding of entorhinal grid-like codes during observation [6-fold symmetrical model, mean grid magnitude (arbitrary units, a.u.) ± s.e.m., 0.124 ± 0.05, one-sample *t*-test, *N* = 45 (one participant excluded due to strategy, one outlier excluded), *t*(44) = 2.257, *d* = 0.34, 95% CI = [0.01, 0.24], *p*_one-tailed_ = 0.015].

## Supplementary Figures

**Figure S1:**
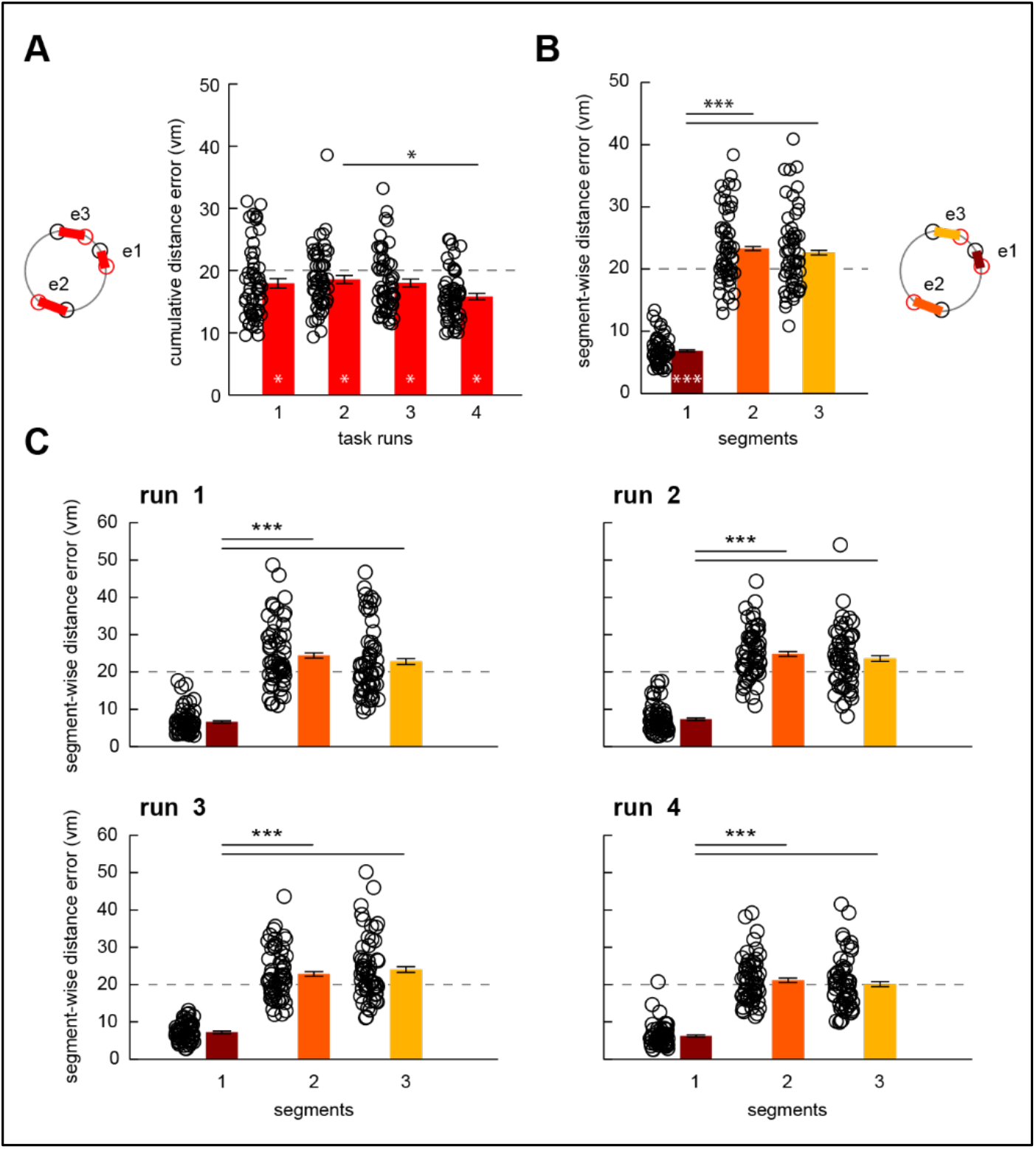
Navigation performance. **(A)** Performance was quantified as cumulative distance error in virtual meters (vm) averaged across all three path errors [e1-e3; showing the difference between the demonstrator’s (black) and participant’s (red) path segment endpoints], indicating the deviation from the demonstrator’s paths. For each of the four task runs, participants reached an average cumulative distance error (virtual meters, vm) significantly below the feedback threshold of 20 vm (dashed line; separate one-sample *t*-tests, all *p*_two-tailed_ < 0.05), whereby the cumulative distance error during the second run was higher compared to the last run [one-way analysis of variance (ANOVA), *N* = 58, main effect of run, *F*(3,227) = 3.53, *η*^*2*^= 0.04, 95% CI = [0, 0.1], *p*_two-tailed_ = 0.016; post-hoc pair-wise comparison run 2 vs. 4, *t*(227) = 3.02, *d* = 0.62, 95% CI = [-5.1, -0.4], *p*_two-tailed_ = 0.015; all other post-hoc comparisons *p*_two-tailed_ > 0.05]. **(B)** Across runs, participants performed best when re-tracing the first path segment [one-way ANOVA, *N* = 58, main effect of path segment, *F*(2,171) = 190.7, *η*^*2*^= 0.69, 95% CI = [0.63, 0.74], *p*_two-tailed_ < 0.0001; pair-wise comparisons: first vs. second segment, *t*(171) = -17.24, *d* = -2.96, 95% CI = [14.2, 18.7], *p*_two-tailed_ < 0.0001, first vs. third segment, *t*(171) = -16.57, *d* = -2.91, 95% CI = [13.5, 18], *p*_two-tailed_ < 0.0001, second vs. third segment, *p*_two-tailed_ = 0.784], significantly below the feedback threshold of 20 vm [separate one-sample *t*-tests, *N* = 58; first segment, *t*(57) = -45.18, *d* = -5.9, 95% CI = [6.3, 7.4], *p*_two-tailed_ < 0.0001; significantly above the feedback threshold: second segment, *t*(57) = 4.3, *d* = 0.56, 95% CI = [21.7, 24.8], *p*_two-tailed_ < 0.0001 ; third segment, *t*(57) = 3.16, *d* = 0.42, 95% CI = [21, 24.3], *p*_two-tailed_ < 0.05]. **(C)** Segment-wise distance errors per run. * *p* < 0.05; *** *p* < 0.0001; error bars reflect the standard error of the mean, s.e.m.

**Figure S2.**
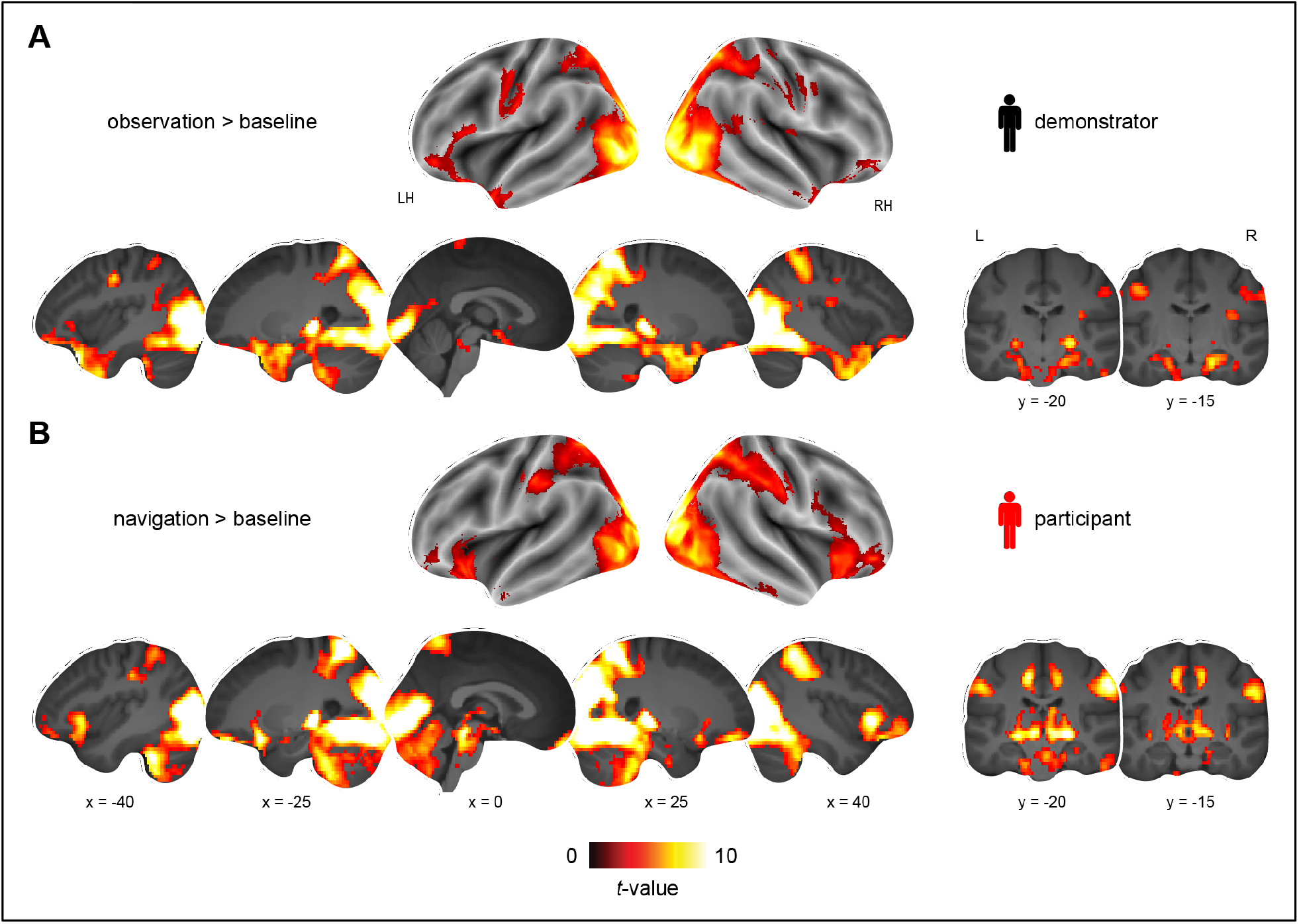
Brain activation profiles during observation and navigation. Brain activation **(A)** during observation and **(B)** navigation, each compared to the implicit (fixation) baseline (**Table S1**), plotted onto brain slices of the average structural image. All results are shown at *p* < 0.05 FWE-corrected at cluster level (cluster-defining threshold of *p* < 0.001).

**Figure S3.**
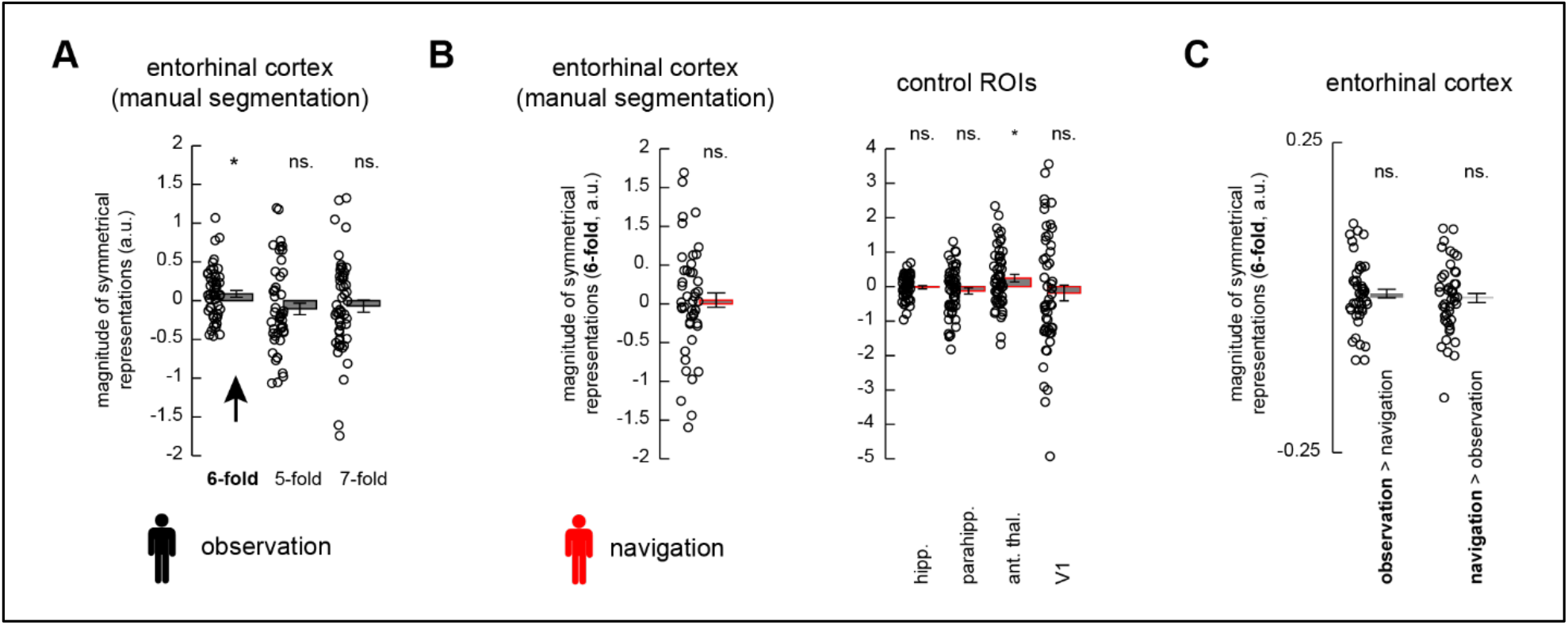
Grid-like codes during observation and navigation. **(A)** Results when manually segmenting the entorhinal cortex ROI. Magnitude of symmetrical representations (6-, 5-, and 7-fold periodicities) in the entorhinal cortex during observation periods (arbitrary units, a.u.). **(B)** No significant entorhinal grid-like codes during navigation periods (again, manual delineation of ROI), and no significant grid-like codes in most of the control ROIs. Notably, grid magnitudes were significantly increased in the anterior thalamus. **(C)** No significant grid-like codes when estimating grid orientations on observation and testing them on navigation periods (and vice versa). * *p* < 0.05; error bars reflect the standard error of the mean, s.e.m.

**Figure S4.**
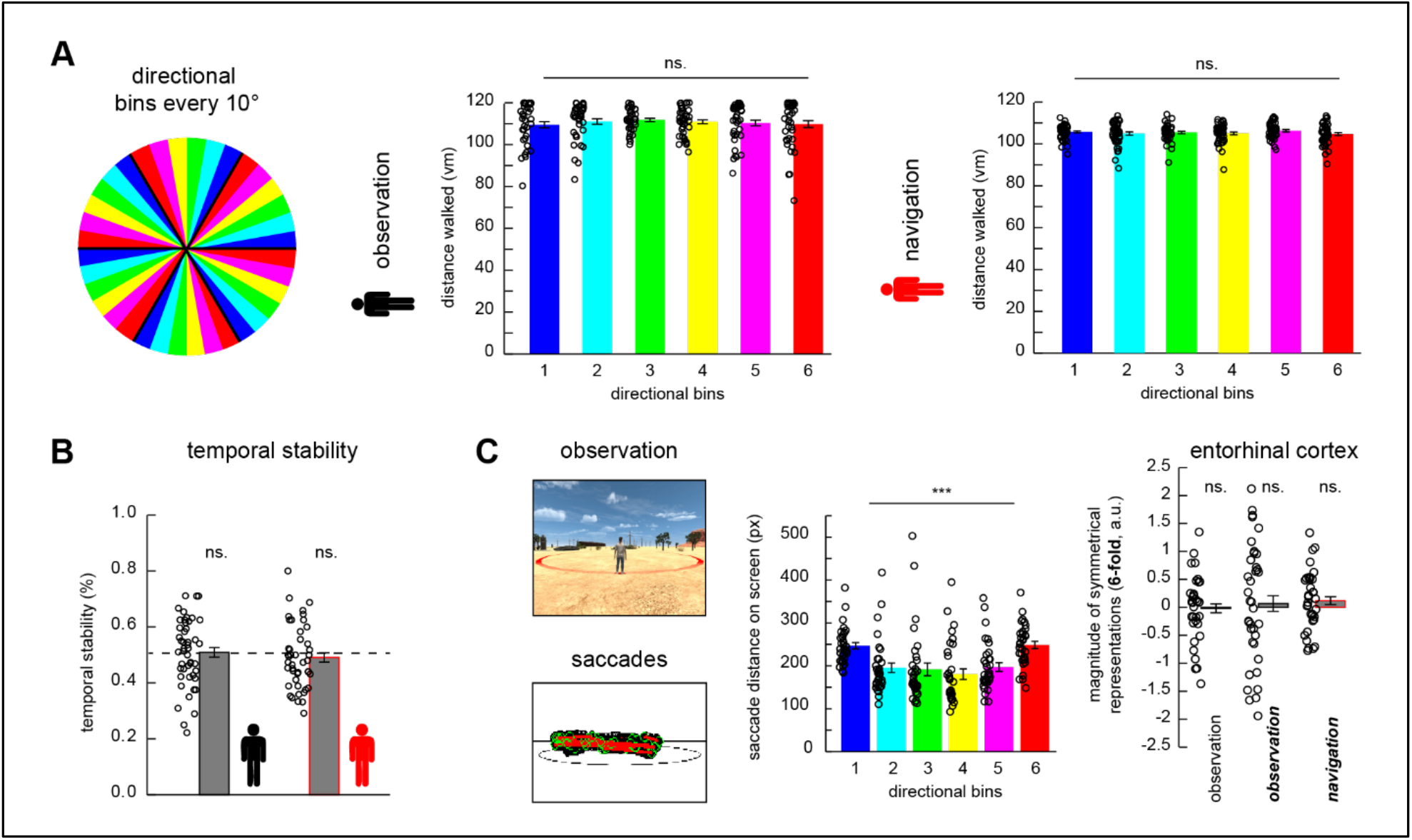
Control analyses. **(A, left panel)** Unbiased estimation of grid orientations. We designed our task such that path segments were oriented along 36 directions that could be divided into 6 directional bins, spanning the 360°-VR-space with 10° angular resolution. **(A, middle panel)** There was no significant difference in the distance walked (vm) across directional bins during observation periods [one-way ANOVA, *N* = 48 (individual data points indicate the trial-wise distance walked in each of the directional bins and in each of the four task runs, based on the two trial sequences randomly generated before the start of the experiment, **Materials and Methods**), no significant main effect of directional bin, *p*_two-tailed_ = 0.896]. **(A, right panel)** This was also not the case during navigation periods [repeated measures ANOVA, *N* = 45 (same participant sample as during initial grid code analysis), no significant main effect of directional bin, *p*_two-tailed_ = 0.968, individual data points indicate the participant-specific distance walked in each of the directional bins, averaged across trials and runs]. **(B)** Temporal stability of grid-like coding was not significantly different between the conditions [paired-sample *t*-test, *N* = 44 (participant sample from which both observation- and navigation-based grid values were available), *p*_two-tailed_ = 0.252] but was generally low (i.e., not significantly different from the temporal stability threshold of 50% for observation/navigation periods marked with dashed line; separate one-sample *t*-tests, *N* = 44, *p*_two-tailed_ = 0.723/0.36), indicating variability of both observation- and navigation-based grid orientations over time. **(C)** No effects of eye movements on entorhinal grid-like coding. There was a significant difference in the saccade distance on the screen across the six directional bins [repeated-measures ANOVA, *N* = 33, *** *p* < 0.0001, individual data points indicate the segment-specific saccade distance on screen (pixels, px) in each of the directional bins, averaged across trials and runs, **Results S4**]. We thus repeated the analysis and randomly selected saccades to match the average, individual saccade distance along directional bins 2 to 5 (analyses marked in bold-italic). Error bars reflect the standard error of the mean, s.e.m; ns., not significant.

**Figure S5.**
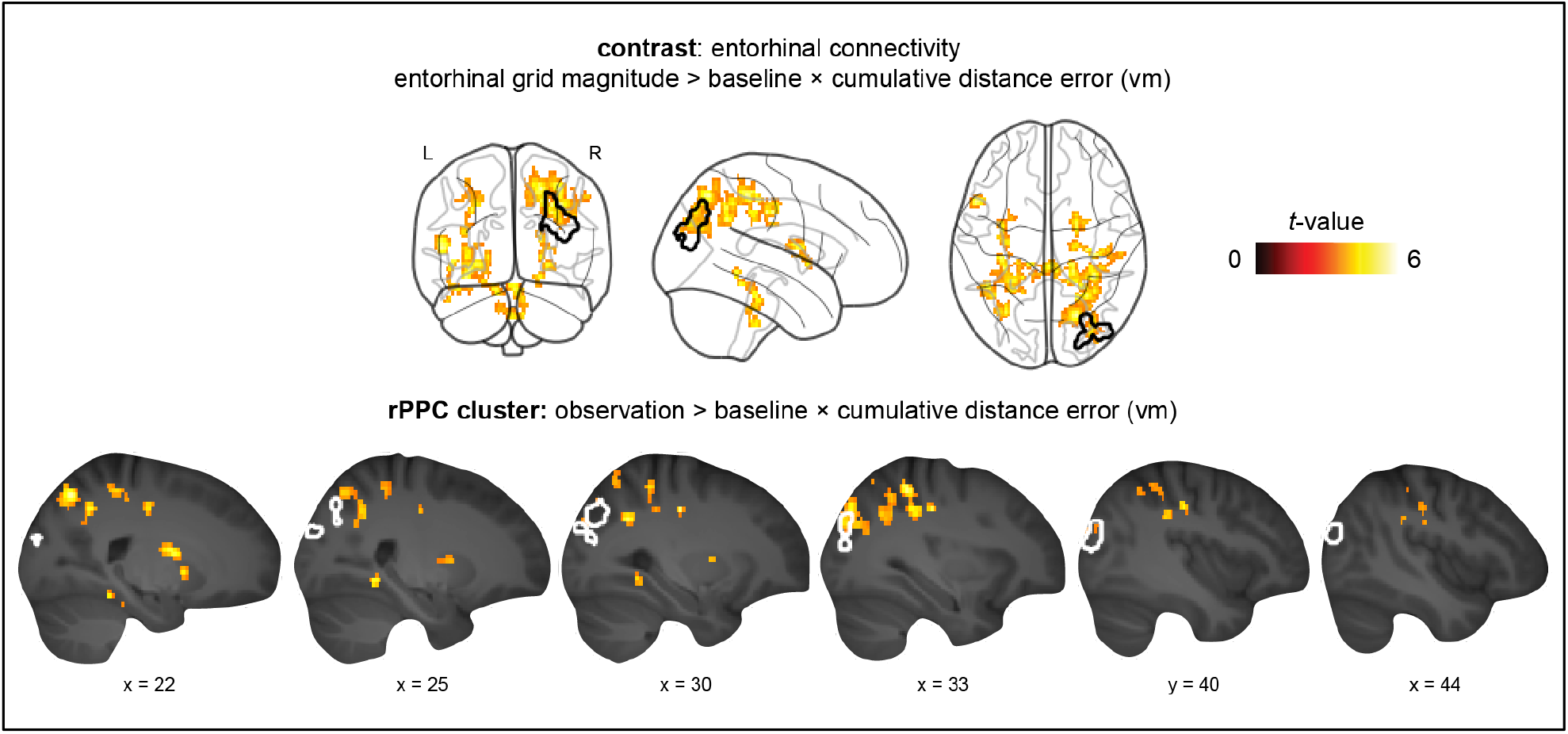
Right posterior parietal cortex contributions during observation. Spatial overlap in the right posterior parietal cortex (rPPC) between entorhinal connectivity × performance results (i.e., entorhinal-cortical coupling during observation, time-locked to grid-like codes and negative association with performance; indicated with warm colors; taken from **Figure 4D**) and activation × performance results (i.e., univariate activity, independent of grid-like codes, and positive association with performance; indicated through black outline; taken from **Figure 2C**).

**Figure S6.**
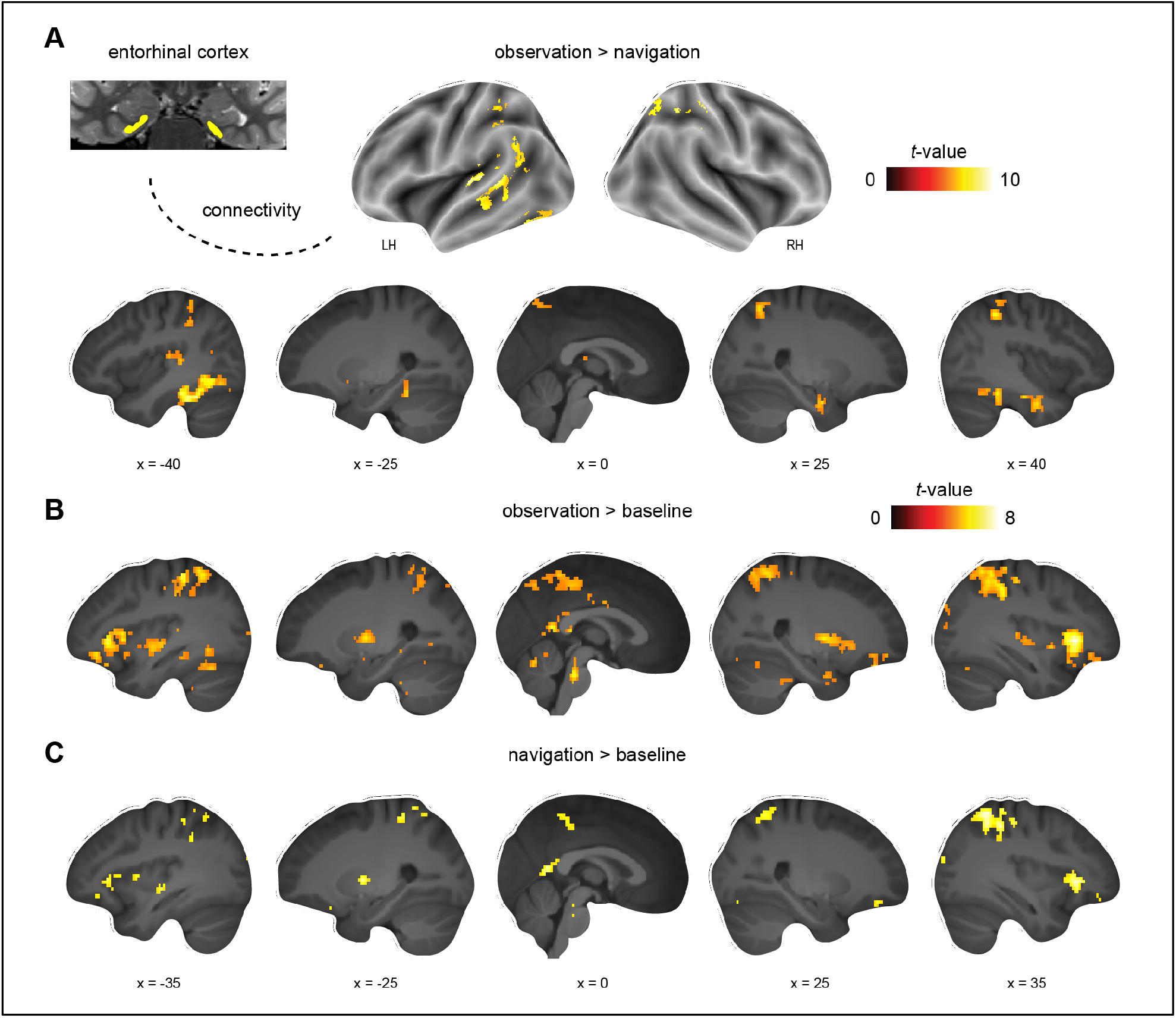
Entorhinal-cortical connectivity changes independent of fluctuations in grid magnitude and performance. **(A)** Increased connectivity during observation compared to navigation. Results are shown at *p* < 0.05 FWE-corrected at cluster level (cluster-defining threshold of *p* < 0.001). **(B-C)** Connectivity changes during observation/navigation compared to the implicit (fixation) baseline. Please note that these results were thresholded at *p* < 0.05 FWE-corrected (i.e., without applying a cluster-defining threshold).

## Supplementary Tables

### General note regarding tables displaying fMRI results

Unless stated otherwise, significance for all MRI analyses was assessed using cluster-inference with a cluster-defining threshold of *p* < 0.001 and a cluster-probability of *p* < 0.05 family-wise error (FWE) corrected for multiple comparisons. The corrected cluster size (i.e., the spatial extent of a cluster that is required in order to be labeled as significant) was calculated using the SPM extension “CorrClusTh.m” and the Newton-Raphson search method (script provided by Thomas Nichols, University of Warwick, United Kingdom, and Marko Wilke, University of Tübingen, Germany)^2^.

MNI coordinates represent the location of cluster peak voxels. We report the first local maximum within each cluster. Complete tables (i.e., also providing the different local maxima within each cluster, reflecting the original output from the abovementioned script to calculate the corrected cluster size) are provided on the OSF^3^.

Anatomical nomenclature for all tables was obtained from the Laboratory for Neuro Imaging (LONI) Brain Atlas (LBPA40, http://www.loni.usc.edu/atlases/; Shattuck et al., 2008). L, left; R, right; LH, left hemisphere; RH, right hemisphere.

**Table S1.** Brain activation profiles during observation and navigation. Analysis consisted of separate one-sample *t*-tests (all *N* = 58), contrasts: observation > navigation (critical cluster size: 91 voxels), navigation > observation (91 voxels), observation > baseline (80 voxels), navigation > baseline (77 voxels).

| Contrast & brain region | MNI |  |  | Z value | Cluster size |
| --- | --- | --- | --- | --- | --- |
|  | x | y | z |  |  |
| **Observation > navigation:** |  |  |  |  |  |
| R middle occipital gyrus | 44 | -82 | -6 | Inf | 7018 |
| L superior temporal gyrus | -48 | -42 | 6 | Inf | 7921 |
| R postcentral gyrus | 36 | -18 | 24 | 7.62 | 1015 |
| R middle frontal gyrus | 18 | 38 | 0 | 7.51 | 3045 |
| L precentral gyrus | -44 | -10 | 39 | 6.75 | 853 |
| L precuneus | -2 | -60 | 45 | 5.97 | 296 |
| L inferior frontal gyrus | -56 | 26 | 0 | 4.98 | 255 |
| **Navigation > observation:** |  |  |  |  |  |
| R superior occipital gyrus | 18 | -88 | 21 | Inf | 39636 |
| R supramarginal gyrus | 56 | -36 | 39 | Inf | 1810 |
| R middle temporal gyrus | 52 | -30 | -9 | 4.49 | 333 |
| **Observation > baseline:** |  |  |  |  |  |
| R middle occipital gyrus | 44 | -80 | -6 | Inf | 33139 |
| L postcentral gyrus | -64 | -6 | 30 | 7.08 | 625 |
| R postcentral gyrus | 68 | -4 | 24 | 5.97 | 642 |
| L postcentral gyrus | -6 | -40 | 66 | 5.53 | 361 |
| R insular cortex | 34 | -18 | 18 | 5.2 | 85 |
| **Navigation > baseline:** |  |  |  |  |  |
| L middle occipital gyrus | -26 | -90 | 18 | Inf | 49749 |
| L middle temporal gyrus | -58 | 6 | -24 | 4.82 | 158 |

**Table S2:** Grid-like codes during observation (results based on the full sample) Results of the main grid code analysis based on two entorhinal cortex ROIs (defined through ASHS and manually segmented using ITK-SNAP) and four control ROIs (hippocampus, parahippocampal cortex, anterior thalamus, and primary visual cortex/V1). *N* describes the full participant sample available for the analyses (i.e., after applying restrictions for voxel selection but no outliers excluded, **Materials and Methods**). Results were corrected for multiple comparisons using Bonferroni-correction (α_Bonferroni_ = 0.05/6 ROIs = 0.008). * *p* < 0.001, (*) *p* < 0.05 but not surviving Bonferroni-correction.

| *Grid-cell-representations during observation* (analysis type) | *N* | Grid magnitude, arbitrary units<br>(mean ± s.e.m.) | *t*-statistic | *p*-value (one-tailed) |
| --- | --- | --- | --- | --- |
| Entorhinal cortex (ASHS), 6-fold rotational symmetry | 49 | 0.157 ± 0.06 | $t(48) = 2.45$ | 0.009 (*) |
| Entorhinal cortex (ASHS), 5-fold rotational symmetry | 49 |  |  | 0.146 |
| Entorhinal cortex (ASHS), 7-fold rotational symmetry | 49 |  |  | 0.23 |
| Entorhinal cortex (ITK-SNAP), 6-fold rotational symmetry | 51 | 0.088 ± 0.05 | $t(50) = 1.9$ | 0.032 (*) |
| Entorhinal cortex (ITK-SNAP), 5-fold rotational symmetry | 51 |  |  | 0.293 |
| Entorhinal cortex (ITK-SNAP), 7-fold rotational symmetry | 51 |  |  | 0.103 |
| Hippocampus, 6-fold rotational symmetry | 58 |  |  | 0.283 |
| Parahippocampal cortex, 6-fold rotational symmetry | 58 |  |  | 0.196 |
| Anterior thalamus, 6-fold rotational symmetry | 58 |  |  | 0.241 |
| Primary visual cortex / V1, 6-fold rotational symmetry | 58 | -0.076 ± 0.17 | $t(57) = -4.462$ | < 0.001 * |

**Table S3:** Grid-like codes during navigation (results based on the full sample) Results of the main grid code analysis based on two entorhinal cortex ROIs (defined through ASHS and manually segmented using ITK-SNAP) and four control ROIs (hippocampus, parahippocampal cortex, anterior thalamus, and primary visual cortex/V1). *N* describes the full participant sample available for the analyses (i.e., after applying restrictions for voxel selection but no outliers excluded, **Materials and Methods**). Results were corrected for multiple comparisons using Bonferroni-correction (α_Bonferroni_ = 0.05/6 ROIs = 0.008). (*) *p* < 0.05 but not surviving Bonferroni-correction, (∼) tendency for *p* < 0.05.

| *Grid-cell-representations during navigation* (analysis type) | *N* | Grid magnitude, arbitrary units<br>(mean ± s.e.m.) | *t*-statistic | *p*-value (one-tailed) |
| --- | --- | --- | --- | --- |
| Entorhinal cortex (ASHS), 6-fold rotational symmetry | 49 | 0.454 ± 0.22 | $t(48) = 1.96$ | 0.028 (*) |
| Entorhinal cortex (ASHS), 5-fold rotational symmetry | 49 |  |  | 0.362 |
| Entorhinal cortex (ASHS), 7-fold rotational symmetry | 49 |  |  | 0.218 |
| Entorhinal cortex (ITK-SNAP), 6-fold rotational symmetry | 51 |  |  | 0.191 |
| Entorhinal cortex (ITK-SNAP), 5-fold rotational symmetry | 51 |  |  | 0.068 (~) |
| Entorhinal cortex (ITK-SNAP), 7-fold rotational symmetry | 51 |  |  | 0.088 |
| Hippocampus, 6-fold rotational symmetry | 58 |  |  | 0.166 |
| Parahippocampal cortex, 6-fold rotational symmetry | 58 |  |  | 0.162 |
| Anterior thalamus, 6-fold rotational symmetry | 58 |  |  | 0.118 |
| Primary visual cortex / V1, 6-fold rotational symmetry | 58 |  |  | 0.118 |

**Table S4:** Entorhinal connectivity changes time-locked to grid-like codes during observation, and association with performance. Analysis consisted of separate one-sample *t*-tests (all *N* = 47, same participant sample as during initial grid code analysis), contrasts: entorhinal grid magnitude during observation > baseline × cumulative distance error across all paths added as a covariate of interest (critical cluster size: 61 voxels). The findings were specific to lower performance (there were no significant connectivity changes related to increased navigation performance) and to observation periods only (there were no significant effects during navigation).

| Contrast & brain region | MNI |  |  | Z value | Cluster size |
| --- | --- | --- | --- | --- | --- |
|  | x | y | z |  |  |
| ***Entorhinal grid magnitude during observation > baseline × covariate cumulative distance error across all paths:*** |  |  |  |  |  |
| R superior parietal gyrus | 22 | -68 | 51 | 4.77 | 613 |
| L inferior frontal gyrus | -52 | 16 | 6 | 4.72 | 73 |
| R postcentral gyrus | 36 | -24 | 39 | 4.61 | 131 |
| L hippocampus | -32 | -34 | -9 | 4.54 | 197 |
| L angular gyrus | -30 | -66 | 36 | 4.34 | 84 |
| Brainstem | 2 | -32 | -24 | 4.31 | 154 |
| R putamen | 22 | 0 | 12 | 4.31 | 92 |
| L middle temporal gyrus | -42 | -48 | -3 | 4.26 | 100 |
| R lingual gyrus | 26 | -48 | -6 | 4.19 | 64 |
| R middle occipital gyrus | 34 | -76 | 39 | 4.09 | 156 |
| L insular cortex | -34 | -6 | 0 | 4.05 | 107 |
| L superior parietal gyrus | -26 | -48 | 39 | 3.98 | 83 |

**Table S5:**
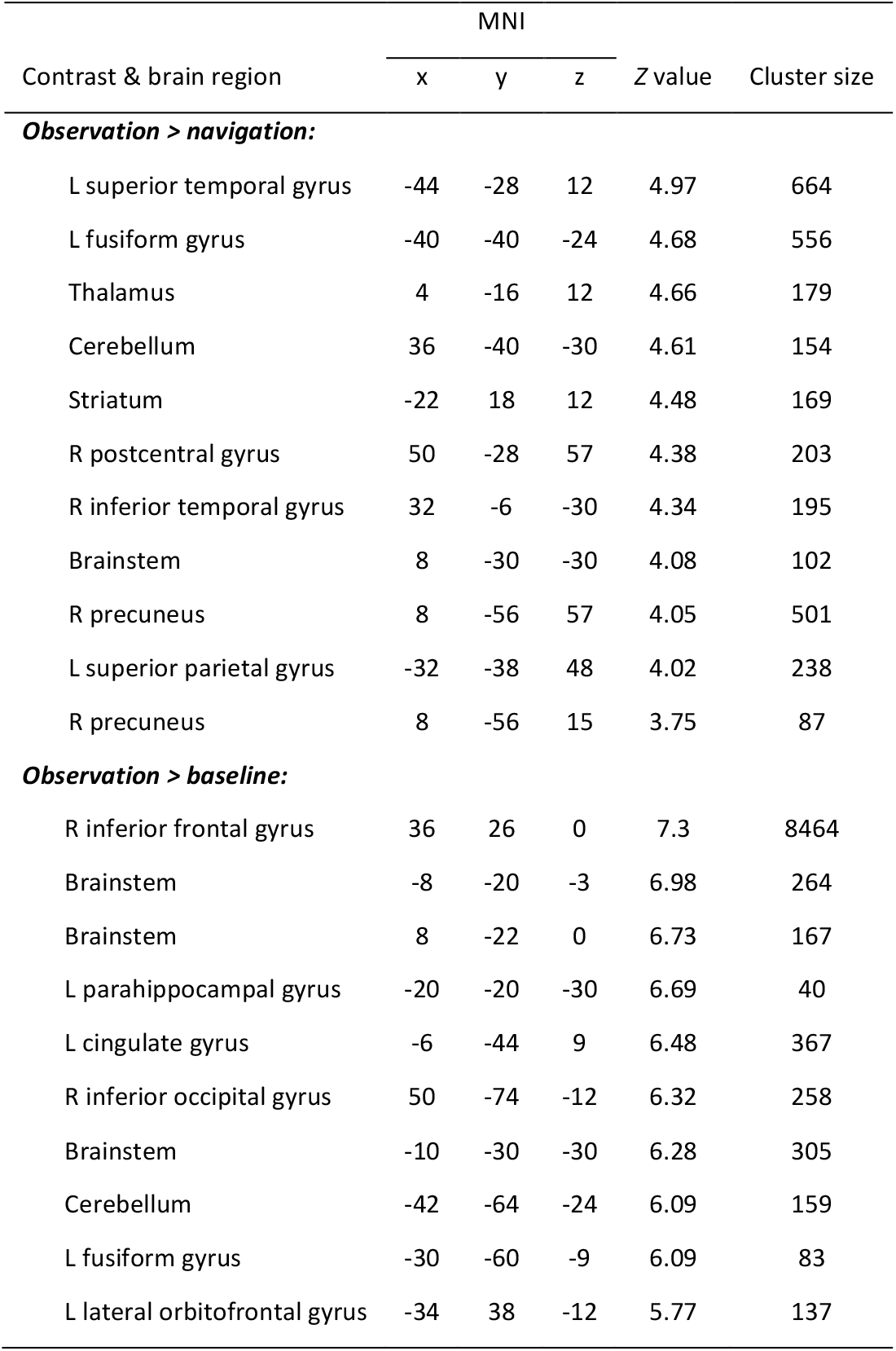
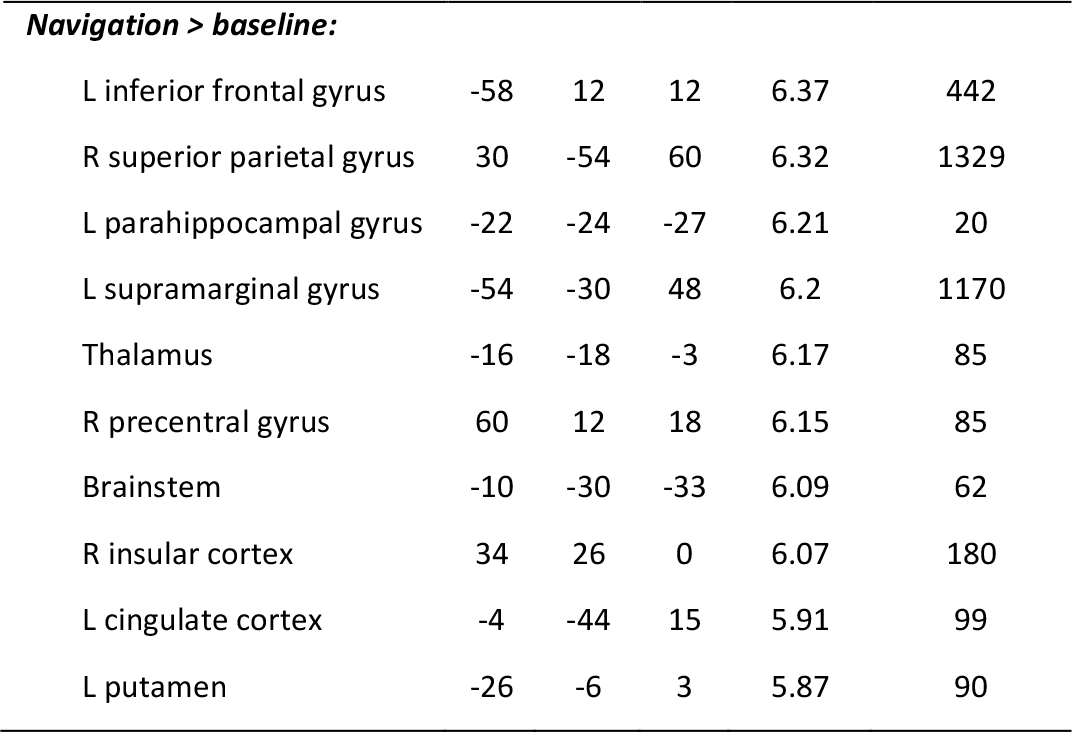
Entorhinal connectivity changes independent of grid-like codes or performance. Analysis consisted of separate one-sample *t*-tests (all *N* = 47, same participant sample as during initial grid analysis), contrasts: observation > navigation (64 voxels), observation > baseline, navigation > baseline (both thresholded using *p* < 0.05 FWE-corrected for multiple comparisons, we report the first 10 clusters). There were no significant results for the reverse contrast navigation > observation.

| Contrast & brain region | MNI |  |  | Z value | Cluster size |
| --- | --- | --- | --- | --- | --- |
|  | x | y | z |  |  |
| **Observation > navigation:** |  |  |  |  |  |
| L superior temporal gyrus | -44 | -28 | 12 | 4.97 | 664 |
| L fusiform gyrus | -40 | -40 | -24 | 4.68 | 556 |
| Thalamus | 4 | -16 | 12 | 4.66 | 179 |
| Cerebellum | 36 | -40 | -30 | 4.61 | 154 |
| Striatum | -22 | 18 | 12 | 4.48 | 169 |
| R postcentral gyrus | 50 | -28 | 57 | 4.38 | 203 |
| R inferior temporal gyrus | 32 | -6 | -30 | 4.34 | 195 |
| Brainstem | 8 | -30 | -30 | 4.08 | 102 |
| R precuneus | 8 | -56 | 57 | 4.05 | 501 |
| L superior parietal gyrus | -32 | -38 | 48 | 4.02 | 238 |
| R precuneus | 8 | -56 | 15 | 3.75 | 87 |
| **Observation > baseline:** |  |  |  |  |  |
| R inferior frontal gyrus | 36 | 26 | 0 | 7.3 | 8464 |
| Brainstem | -8 | -20 | -3 | 6.98 | 264 |
| Brainstem | 8 | -22 | 0 | 6.73 | 167 |
| L parahippocampal gyrus | -20 | -20 | -30 | 6.69 | 40 |
| L cingulate gyrus | -6 | -44 | 9 | 6.48 | 367 |
| R inferior occipital gyrus | 50 | -74 | -12 | 6.32 | 258 |
| Brainstem | -10 | -30 | -30 | 6.28 | 305 |
| Cerebellum | -42 | -64 | -24 | 6.09 | 159 |
| L fusiform gyrus | -30 | -60 | -9 | 6.09 | 83 |
| L lateral orbitofrontal gyrus | -34 | 38 | -12 | 5.77 | 137 |
|  | x | y | z |  |  |
| ***Navigation > baseline:*** |  |  |  |  |  |
| L inferior frontal gyrus | -58 | 12 | 12 | 6.37 | 442 |
| R superior parietal gyrus | 30 | -54 | 60 | 6.32 | 1329 |
| L parahippocampal gyrus | -22 | -24 | -27 | 6.21 | 20 |
| L supramarginal gyrus | -54 | -30 | 48 | 6.2 | 1170 |
| Thalamus | -16 | -18 | -3 | 6.17 | 85 |
| R precentral gyrus | 60 | 12 | 18 | 6.15 | 85 |
| Brainstem | -10 | -30 | -33 | 6.09 | 62 |
| R insular cortex | 34 | 26 | 0 | 6.07 | 180 |
| L cingulate cortex | -4 | -44 | 15 | 5.91 | 99 |
| L putamen | -26 | -6 | 3 | 5.87 | 90 |

## Footnotes

1 To clarify, we assume that the majority of participants formed a “mental map”, that is, a sense of the general layout of the environment as it would have been very difficult to successfully perform the task otherwise. Unfortunately, we do not have more detailed data available on these participants as no follow-up questions were asked about how these participants actually created their “mental maps”.

2 http://www2.warwick.ac.uk/fac/sci/statistics/staff/academic-research/nichols/scripts/spm/

3 https://osf.io/mhtgp/?view_only=ff7038b65c0345cd9ffa4ccd3813d7ba, *note that the OSF project website is currently set to private and will be made publicly available upon publication of the manuscript*

